# Nature-inspired nanodiscs for lesion-targeted delivery reprogram macrophages and attenuate established abdominal aortic aneurysms

**DOI:** 10.64898/2026.05.13.724870

**Authors:** Zhongshan He, Yitao Huang, Yinghui Wang, Qian Ren, Junchao Xu, Qiaochu Wang, Lian-Wang Guo, Huan Bao

**Affiliations:** Department of Molecular Physiology and Biological Physics, University of Virginia, Charlottesville, VA 22903, USA; Division of Surgical Sciences, Department of Surgery, School of Medicine, University of Virginia, Charlottesville, VA 22903, USA; Department of Bioengineering, University of Pennsylvania, Philadelphia, PA 19104, USA

**Author notes:** Corresponding authors: L.G., H.B. These authors contributed equally: Zhongshan He, Yitao Huang.

**Keywords:** Abdominal aortic aneurysms, Macrophage reprogramming, Nanodiscs, Immunomodulation, Targeted delivery

## Abstract

Abdominal aortic aneurysm (AAA) is a life-threatening vascular disease characterized by chronic inflammation and immune dysregulation, with lesional macrophages playing a pivotal role in disease progression. However, effective and safe delivery of immune modulators to macrophages at the site of AAA remains a major clinical challenge. To address this unmet need, we report a nature-inspired nanodisc platform based on high-density lipoproteins for targeted delivery to lesional macrophages, further engineered with a multi-component targeting strategy incorporating an aneurysm-homing peptide and phosphatidylserine lipids. Nanodiscs encapsulating an anti-inflammatory protein kinase R-like endoplasmic reticulum kinase (PERK) inhibitor remarkably attenuated progression of established AAA in an elastase-induced mouse model. Using a combination of in vivo biodistribution and immune profiling approaches, we demonstrate that nanodisc-assisted PERK inhibitor delivery selectively reprograms the local immune microenvironment and attenuates pathological inflammation in AAA disease models. Notably, a single administration achieves sustained therapeutic efficacy with favorable safety profiles, effectively limiting the progression of established AAA in a clinically relevant setting. This work presents a new avenue of designer nanomedicines for targeted immunomodulation and maybe broadly applicable for a wide range of vascular and immune-mediated pathologies.

## Introduction

Abdominal aortic aneurysm (AAA) is a progressive and life-threatening inflammatory vascular disease characterized by permanent dilation of the abdominal aorta and a high risk of rupture, leading to substantial morbidity and mortality, particularly in the elderly population.^1–2^ Most patients with AAA remain asymptomatic until catastrophic rupture, which is frequently fatal and poses major challenges for early diagnosis and clinical management.^3^ Current clinical guidelines recommend open surgical repair or endovascular aneurysm repair primarily for large, symptomatic, or ruptured aneurysms;^4–5^ however, these interventions are invasive, associated with perioperative complications, and show limited long-term durability, with approximately 20-30% of patients requiring reintervention within five years after endovascular repair.^6–8^ Consequently, patients with small or moderate asymptomatic aneurysms, or those unsuitable for surgery due to anatomical or clinical constraints, are typically managed by surveillance alone, leaving disease progression unaddressed and imposing substantial clinical and psychological burdens.^8^ Despite advances in surgical and imaging technologies, there are currently no US Food and Drug Administration-approved medical therapies to block AAA progression or reduce the risk of rupture.^9–10^ Given that chronic inflammation and immune dysregulation are central drivers of AAA pathogenesis,^2^ precision therapeutic strategies that selectively neutralize these processes within aneurysmal lesions represent a compelling and unmet clinical need.

Accumulating evidence indicates that macrophages play a central role in the initiation and progression of AAA by orchestrating chronic inflammation, excessive reactive oxygen species (ROS) production, and extracellular matrix degradation within aneurysmal lesions.^11–12^ Activated lesional macrophages secrete pro-inflammatory cytokines, chemokines, and proteolytic enzymes, thereby amplifying inflammatory cascades, promoting oxidative stress, and accelerating elastin fragmentation and medial degeneration of the aortic wall.^11–13^ By virtue of their abundance, functional plasticity, and pivotal contribution to disease pathology, macrophages have emerged as an attractive therapeutic target for AAA intervention. In this context, nanomedicine-based delivery strategies have been increasingly explored for AAA treatment owing to their ability to improve drug stability, prolong circulation time, and enable localized therapeutic action.^14–16^ Various nanotherapeutic platforms, including polymeric nanoparticles, lipid-based carriers, and biomimetic systems, have been developed to deliver anti-inflammatory or antioxidative agents to aneurysmal tissues.^17–20^ However, the clinical translation of these nanotherapeutics remains limited by insufficient accumulation at disease sites, complex fabrication processes, and challenges associated with repeated administration and long-term biosafety, thereby limiting their clinical translation.^21–23^ Consequently, there is a pressing challenge to develop biocompatible low-immunogenic delivery platforms that effectively modulate oxidative stress and inflammation, with efficient and selective targeting of macrophage-rich aneurysmal lesionss being key to maximizing therapeutic efficacy and minimizing systemic adverse effects.

Nature-inspired, synthetic high-density lipoprotein–mimicking (sHDL) nanodiscs represent a unique class of biomimetic nanoplatforms that combine excellent biocompatibility with compelling physicochemical properties for inflammatory disease treatment.^24–27^ Owing to their low immunogenicity and structural similarity to endogenous HDL particles, nanodiscs offer a safe and effective delivery system that is well-suited for chronic inflammatory diseases and cardiovascular disorders.^28–32^ Their nanoscale size (<100 nm) and disc-like morphology confer enhanced vascular wall penetration and lesional infiltration, thereby facilitating efficient uptake by deeply embedded target cells within the lesion, such as lesional macrophages.^33–36^ Importantly, sHDLs have previously been manufactured for clinical testing and demonstrated favorable safety profiles in humans, with a maximum tolerated dose of approximately 2.2 g/m^2,37–39^ which is one to two orders of magnitude higher than that reported for most polymeric or inorganic nanoparticles evaluated in clinical trials.^40^ These properties underscore the remarkable translational potential of sHDL-based platforms. Nevertheless, the utility of sHDL nanodiscs as a targeted nanotherapeutic for AAA, a disease defined by the very inflammatory processes these platforms are equipped to address, remains unexplored.

To bridge this gap, we rationally engineered sHDL nanodiscs with a multi-component targeting strategy by incorporating phosphatidylserine (PS) lipids and the S2P peptide developed specifically to bind the scavenger receptor stabilin-2,^41–45^ which together enable precise recognition of macrophage-rich aneurysmal lesions. This targeting strategy is designed to overcome the limitations of passive accumulation, thereby maximizing local delivery efficiency while minimizing off-target distribution. Moreover, we loaded an inhibitor for PERK (PERKi), identified in our previous studies as a promising therapeutic target for AAA^46^, into the sHDL nanodiscs to selectively modulate inflammatory and immune activation within lesional macrophages.^47–49^ In doing so, our sHDL-based nanoplatform achieves targeted delivery for immune reprogramming of lesional macrophages to resolve inflammation. By combining biomimetic engineering, macrophage targeting, and modulation of PERK-mediated inflammatory pathways, this work establishes a versatile nanoplatform for chronic inflammatory vascular diseases (**Fig.1**). Notably, a single administration of our nanotherapeutic enables effective control of established AAA, underscoring a clinically relevant therapeutic paradigm that remains underexplored in this disease context and may offer improved patient compliance. Collectively, our findings highlight the broad utility of sHDL nanodiscs as a next-generation strategy for targeted immunomodulation and provide a conceptual framework for the development of precision nanomedicines in AAA and other immune-mediated pathologies.

**Fig. 1.**
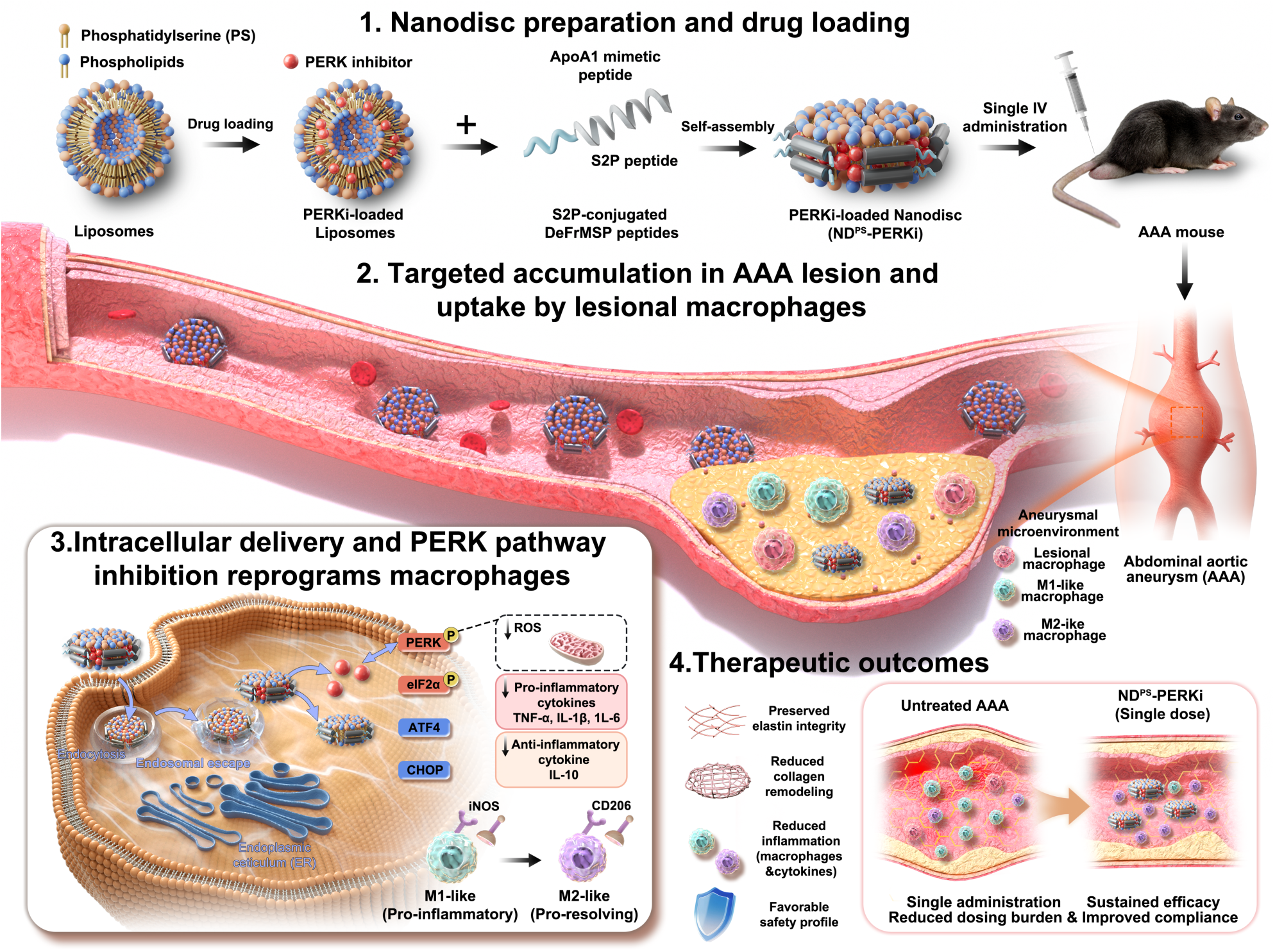
Schematic illustration of the design, targeting strategy, and therapeutic mechanism of engineered nanodiscs for abdominal aortic aneurysm (AAA) treatment. Nanodiscs targeting AAA were fabricated through a self-assembly process using ApoA1 mimetic peptides and phospholipids followed by efficient loading of a PERK inhibitor. The nanodiscs were functionalized with S2P peptide and phosphatidylserine (PS) to establish a dual-targeting system, enabling enhanced homing to AAA lesions and selective uptake by lesional macrophages after systemic administration. Upon accumulation within the aneurysmal microenvironment, engineered nanodiscs are internalized by macrophages and releases the PERK inhibitor intracellularly, leading to inflammatory responses. This process reprograms pro-inflammatory M1-like macrophages toward a pro-resolving M2-like phenotype, characterized by reduced production of pro-inflammatory cytokines and enhanced anti-inflammatory signaling. As a result, nanodisc treatment alleviates vascular inflammation, preserves elastin integrity, and limits pathological remodeling within the aneurysmal wall, ultimately restraining AAA progression. Notably, a single systemic administration achieves sustained therapeutic efficacy with a favorable safety profile, highlighting the translational potential of this nanodisc-based immunomodulatory strategy for AAA and other macrophage-driven vascular inflammatory diseases.

## Results

### Study design

Fig. 1 presents a schematic overview of the design, targeting strategy, and therapeutic mechanism of nanodisc-assisted PERK inhibitor delivery for abdominal aortic aneurysm (AAA) treatment. We first developed and characterized AAA-targeting nanodiscs, including their physicochemical properties, morphology, and structural features. We then investigated their cellular internalization, ROS-scavenging capability, as well as their ability to reprogram macrophage and modulate inflammatory responses in vitro. Subsequently, we assessed the in vivo pharmacokinetics, biodistribution, AAA lesion-targeting capability, and therapeutic efficacy of engineered nanodiscs in an elastase-induced mouse model of AAA. Finally, we elucidated the therapeutic mechanisms of our strategy and evaluated its overall safety profile.

### Constructing sHDL nanodiscs to target inflammatory macrophages

Peptide-encircled sHDL nanodiscs have emerged as promising vehicles for therapeutic delivery, yet many existing peptide scaffolds lack specific targeting modules and are only compatible with DMPC lipids. To address this limitation, we previously developed detergent-free membrane scaffold peptides (DeFrMSPs) that readily transform liposomes of diverse lipid compositions into nanodiscs in a single, detergent-free step (Fig. 2A).^24^ Building on this platform, we sought to determine whether DeFrMSPs could be further functionalized with targeting sequences to enhance cell-specific delivery.

**Fig. 2.**
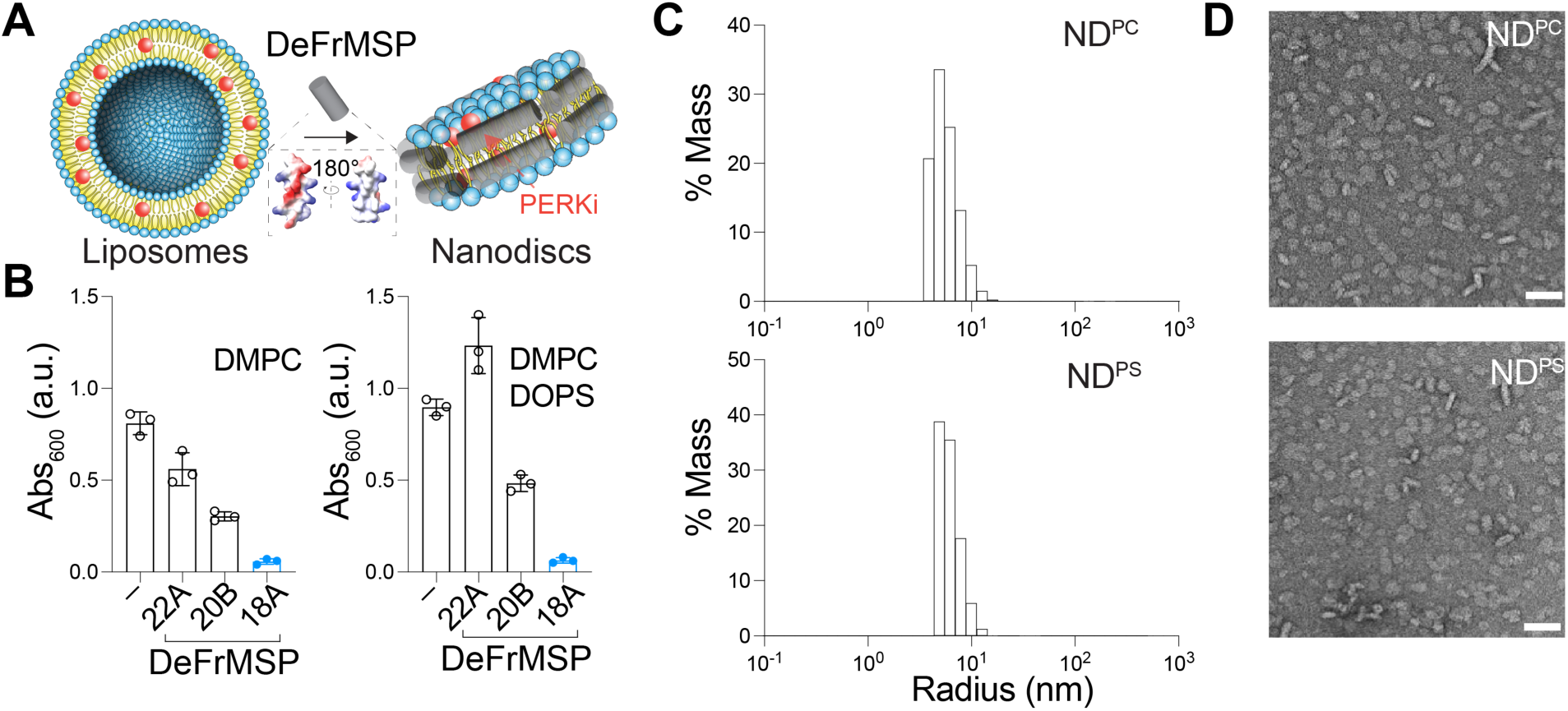
Preparation and characterization of NDs. **(A)** Schematic illustration of the design and preparation workflow of generating sHDL nanodiscs (NDs) for the delivery of a PERK inhibitor (PERKi) using DeFrMSP peptides appended with the S2P targeting sequence. Liposomes encapsulating PERKi are transformed into NDs in one step by incubation with highly amphipathic DeFrMSPs. Inset, structural model of DeFrMSP_18A_ with hydrophobic residues shown in white and hydrophilic residues in red and blue. **(B)** Optical intensities of liposomes prepared with the indicated lipids at 600 nm after incubation with selected DeFrMSP-S2P designs. **(C)** Size distribution of nanodiscs from DLS measurements. Nanodiscs were made using DeFrMSP_18A_-S2P and liposomes harboring DMPC (ND^PC^) or DMPC/DOPS lipids (ND^PS^). **(D)** Representative negative stain EM micrograph of NDs. Scale bar, 30nm. Similar results were obtained with three independent experiments.

To this end, we appended several of our most robust DeFrMSPs with the S2P targeting sequence (CRTLTVRKC), which selectively recognizes inflammatory macrophages. One resulting design, DeFrMSP_18A_-S2P, retained robust nanodisc-forming activities, efficiently converting DMPC liposomes into water-soluble nanodiscs within a few hours of incubation on ice, as evidenced by a marked decrease in optical density at 600 nm (Fig. 2B, left). By contrast, S2P conjugation was poorly tolerated by DeFrMSP_22A_ and DeFrMSP_20B_, with the former showing almost no detectable nanodisc-forming activity and the latter only a modest reduction in liposome absorbance. Leveraging the broad lipid compatibility of DeFrMSPs, we also generated nanodiscs incorporating DOPS lipids, which are preferentially recognized by macrophages relative to other cell types (Fig. 2B, right).^42^ These particles were characterized by dynamic light scattering and negative stain electron microscopy (Fig. 2C and D), confirming the expected discoidal morphology with diameters of approximately 10-20 nm. Thus, DeFrMSP_18A_-S2P nanodiscs formulated with DMPC or DMPC/DOPS lipids were selected for subsequent delivery studies and are referred to hereafter as ND^PC^ and ND^PS^, respectively.

### In vitro comparison of cellular uptake of targeted nanodiscs

Efficient cellular uptake is a critical prerequisite for evaluating the utility of our rationally engineered, macrophage-targeted nanodiscs. Hence, we prepared DiD-labeled ND^PC^ and ND^PS^ to compare their uptake efficiency and determine whether the S2P targeting peptide and DOPS lipids synergistically enhance nanodisc internalization by macrophages. RAW264.7 macrophages were pre-treated with lipopolysaccharide (LPS) to recapitulate the pro-inflammatory microenvironment characteristic of abdominal aortic aneurysm lesions,^44,50–52^ and cellular uptake was subsequently quantified by confocal microscopy (Fig. 3A, B and Fig. S1) and flow cytometry (Fig. 3C and D).

**Fig. 3.**
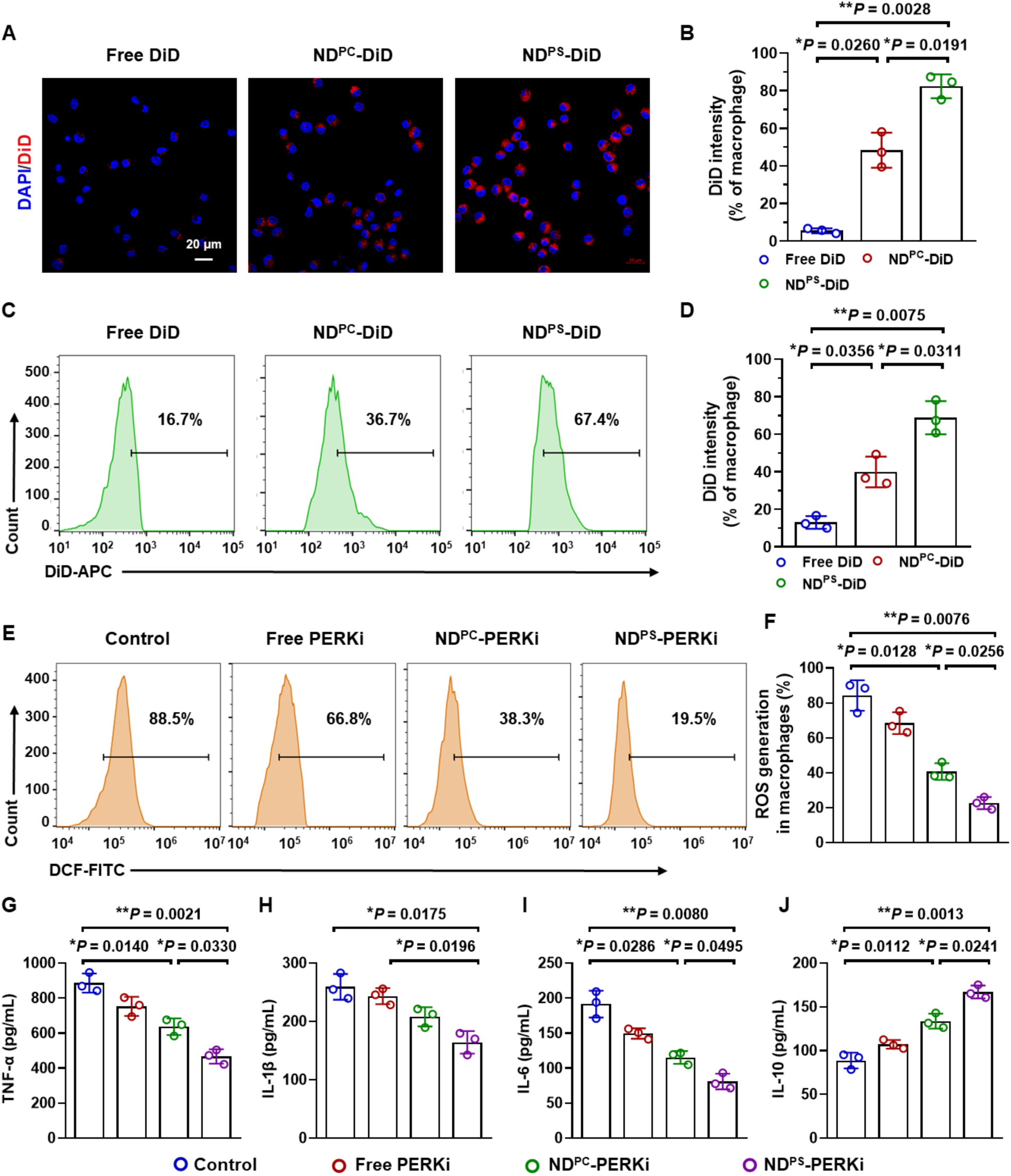
Comparison of cellular uptake, ROS-scavenging capability, and anti-inflammatory efficacy in macrophage after treatment with different targeted nanodiscs. **(A)** Fluorescence microscope images showing the cellular uptake of free DiD and DiD-labeled ND^PC^ or ND^PS^ in LPS-stimulated RAW264.7 macrophages. RAW264.7 cells were incubated in fresh medium containing LPS (500 ng/mL) for 12 hours, followed by treatment with various DiD-labeled formulations for 4 hours. Cellular uptake was visualized using a confocal microscope with 633 nm laser excitation. Red: NDs labeled with DiD; blue: nuclei stained with DAPI. Scale bar = 20 μm. **(B)** Quantitative analysis of DiD fluorescence intensity in macrophage from confocal laser scanning microscope images. **(C-D)** Representative flow cytometry histograms (C) and quantification (D) of cellular uptake of DiD-labeled NDs in LPS-stimulated RAW264.7 macrophages. **(E-F)** Representative flow cytometry histograms (E) and quantification (F) of intracellular ROS generation of RAW264.7 macrophages after different treatments, including free PERK inhibition (PERKi), ND^PC^-PERKi, and ND^PS^-PERKi for 2 h followed by co-incubation with LPS (500 ng/mL) for 12 h. RAW264.7 macrophages treated with LPS only serves as the Control group. **(G-J)** Secretion levels of typical pro-inflammatory cytokines (TNF, IL-1β and IL-6; G-I) and typical anti-inflammatory cytokine IL-10 (J) by RAW264.7 cells were measured by ELISA. Data were analyzed using one-way ANOVA with a Games-Howell post hoc test and shown as mean ± S.D. (n = 3 biologically independent samples). Statistical significance is indicated as*P < 0.05, **P < 0.01, and P > 0.05 denotes no significance.

Both ND^PC^-DiD and ND^PS^-DiD were efficiently internalized by LPS-stimulated RAW264.7 macrophages, while free DiD showed negligible uptake (Fig. 3A-D), demonstrating the ability of nanodiscs to facilitate intracellular delivery of membrane-embedded lipophilic cargo. Notably, ND^PS^-DiD demonstrated substantially greater intracellular accumulation than ND^PC^-DiD, likely reflecting the synergistic contributions of the S2P targeting peptide and PS-mediated macrophage recognition. Since S2P and PS lipids engage distinct cell-surface receptors, these results suggest that the dual-ligand surface design of ND^PS^-DiD greatly enhances macrophage uptake, most likely through receptor-mediated endocytosis.

### Engineered nanodiscs to target the PERK pathway in macrophages

Given the enhanced macrophage uptake observed above and the reactive oxygen species (ROS)-rich inflammatory microenvironment of abdominal aortic aneurysm, we next characterized the ROS-scavenging and anti-inflammatory capabilities of ND^PC^ and ND^PS^ loaded with GSK2606414, a hydrophobic PERK inhibitor (PERKi) with promising therapeutic effects in controlling pre-existing aneurysmal lesions.^46^

To monitor intracellular ROS production, we employed 2’,7’-dichlorofluorescein diacetate (DCFH-DA) staining analyzed by flow cytometry analysis (Fig. 3E and F).^44^ LPS treatment drastically elevated ROS levels in RAW264.7 macrophages, as reflected by high DCF fluorescence intensities (Fig. 3E and F). PERKi treatment considerably reduced intracellular ROS, with the greatest reduction observed when PERKi was delivered via nanodiscs. Notably, ND^PS^-PERKi outperformed ND^PC^-PERKi, consistent with the superior intracellular delivery conferred by the dual targeting moieties (S2P and PS), resulting in more effective ROS scavenging and antioxidative activity.

We next evaluated anti-inflammatory efficacy by measuring AAA-associated cytokine secretion from LPS-stimulated RAW264.7 macrophages^44^, including the pro-inflammatory cytokines tumor necrosis factor-α (TNF-α), interleukin-1β (IL-1β), and interleukin-6 (IL-6), and the anti-inflammatory cytokine interleukin-10 (IL-10). As expected, LPS treatment caused elevated pro-inflammatory cytokine production and reduced IL-10 secretion (Fig. 3G-J). PERKi-loaded nanodiscs effectively suppressed TNF-α, IL-1β, and IL-6 (Fig. 3G-I) while enhancing IL-10 secretion (Fig. 3J), mirroring the trend observed for ROS scavenging^44,52^. Importantly, empty nanodiscs did not exacerbate inflammatory activation under LPS-stimulated conditions (Fig. S2) and instead modestly promoted inflammation resolution, with ND^PS^ exhibiting the most pronounced effect, suggesting intrinsic immunomodulatory properties of the nanodisc platform. The superior anti-inflammatory performance of ND^PS^ reinforced its selection for subsequent in vivo studies and highlights the unique advantage of this delivery platform in inflammatory disease settings. As such, we posit that nanodisc-assisted delivery of PERKi suppresses ROS and pro-inflammatory signaling, providing a compelling mechanistic rationale for evaluating its therapeutic potential in AAA models.

### Enhanced enrichment of targeted nanodiscs in aneurysmal lesion–macrophages

We next characterized the in vivo biodistribution and aneurysmal lesion-targeting efficiency of nanodiscs in AAA and healthy non-AAA mice. DiD-labeled ND^PC^ and ND^PS^ were administered by intravenous injection, and aortas along with major organs (heart, liver, spleen, lung, and kidney) were harvested 24 hours post-injection for ex vivo near-infrared fluorescence (NIRF) imaging (Fig. 4A and Fig. S3).

**Fig. 4.**
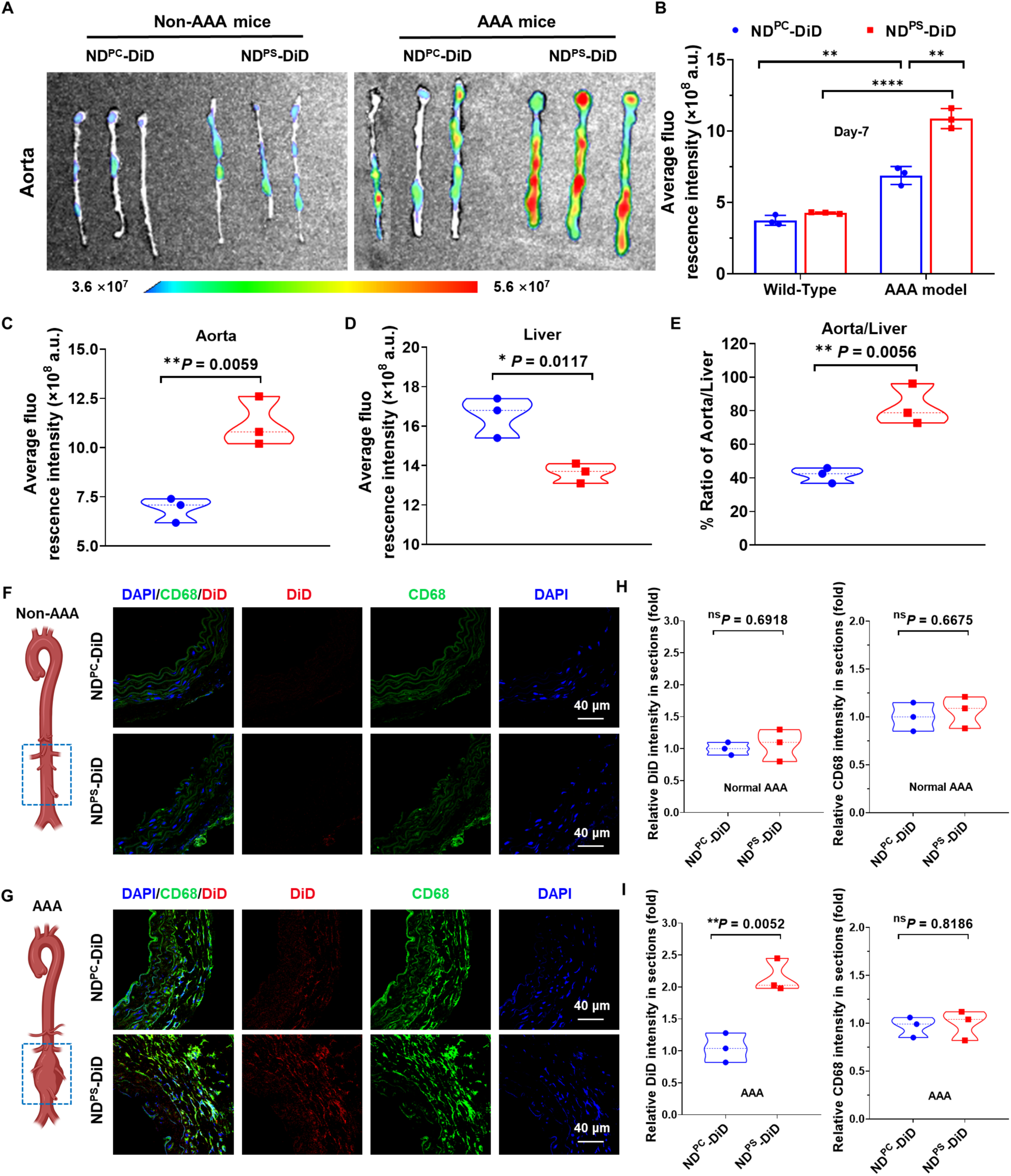
*In vivo* biodistribution and lesional targeting performance of NDs in abdominal aortic aneurysm (AAA) mice. **(A-B)** Representative IVIS images (A) and quantification (B) of DiD fluorescence signals in the aorta of abdominal aortic aneurysm (AAA) mice or C57BL/6J healthy non-AAA mice. The mice were sacrificed 24 h post-intravenous injection of ND^PC^-DiD or ND^PS^-DiD (100 µL, 2 µM DiD per mouse) (n = 3 biologically independent mice). The AAA mice were generated by incubating ∼0.6 U elastase around the abdominal aorta for 8 minutes, followed by a 7-day waiting period before administration of DiD-labelled NDs. **(C-E)** Quantification of the ratio of lesional aorta-to-liver distribution for ND^PC^-DiD and ND^PS^-DiD in AAA mice (n = 3 biologically independent mice). **(F-G)** Confocal microscopy images of abdominal aortic sections from C57BL/6J healthy non-AAA mice (F) or AAA mice (G) treated with ND^PC^-DiD or ND^PS^-DiD. Green: macrophage marker is stained with anti-CD68 antibody; Red: DiD labeled NDs; Blue: DAPI staining for cell nuclei. Scale bars, 40 µm. **(H-I)** Quantification of the fluorescence intensity ratio of DiD and CD68 in the abdominal aortic sections shown in C57BL/6J healthy mice (H) or AAA mice (I) in panel F and G (n = 3 biologically independent mice). Data in (C-E, H, and I) were represented as the violin plots indicating median (middle line), 25th, 75th percentile (violin), as well as minimum and maximum values (whiskers) Data were analysed using unpaired two-tailed Student’s t-test., and shown as mean ± S.D. *P < 0.05, **P < 0.01, ***P < 0.001 and ****P < 0.0001. NS, not significant.

ND^PS^ exhibited greater targeting specificity for aneurysmal aortas compared with ND^PC^, with approximately 1.6-fold higher DiD signal intensity (Fig. 4A and B). In contrast, fluorescence signals detected in the aortas of healthy mice treated with either nanodisc were markedly lower than those observed in AAA mice (Fig. 4B), confirming disease-selective accumulation of targeted nanodiscs in aneurysmal lesions.

Biodistribution analysis further revealed reduced liver uptake of ND^PS^ relative to ND^PC^ (Fig. S3), consistent with its preferential accumulation in aneurysmal tissue. Quantitative analysis of DiD fluorescence in isolated aortas and livers (Fig. 4C and D) found that ND^PS^ achieved an approximately 2-fold higher aorta-to-liver fluorescence ratio than ND^PC^ in AAA mice (Fig. 4E), indicating superior lesion-targeting efficiency and reduced off-target hepatic accumulation.

To further investigate macrophage-specific accumulation within aneurysmal lesions, aortic sections from AAA and healthy non-AAA mice were subjected to immunofluorescence staining for macrophages (CD68) and nuclei (DAPI). As shown in Fig. 4F-I, immunofluorescence analysis revealed much higher DiD fluorescence (DiD, red) co-localized with lesional macrophages (CD68, green) in the abdominal aortas of AAA mice treated with nanodiscs compared with healthy mice.

Notably, DiD signal intensity within lesional areas was enriched by over 2-fold in ND^PS^-treated mice than in ND^PC^-treated mice (Fig. 4G and 4I), indicating more efficient macrophage targeting through the combined contributions of S2P peptide and PS lipid recognition. These findings demonstrate that ND^PS^ effectively penetrates the arterial wall and preferentially accumulates in lesional macrophages, establishing the groundwork for subsequent in vivo therapeutic evaluation of ND^PS^ as a PERKi delivery vehicle in AAA models.

### Nanodisc-assisted delivery of PERKi ameliorates AAA progression

Encouraged by the enhanced aneurysmal lesion and macrophage targeting of ND^PS^, we next evaluated the in vivo therapeutic efficacy of ND^PS^-PERKi in an elastase-induced AAA mouse model (Fig. 5A). AAA mice were randomly assigned to receive intravenous saline, free PERKi, ND^PS^-PERKi, or blank ND^PS^, followed by comprehensive morphological and imaging-based assessments of aneurysm progression.

**Fig. 5.**
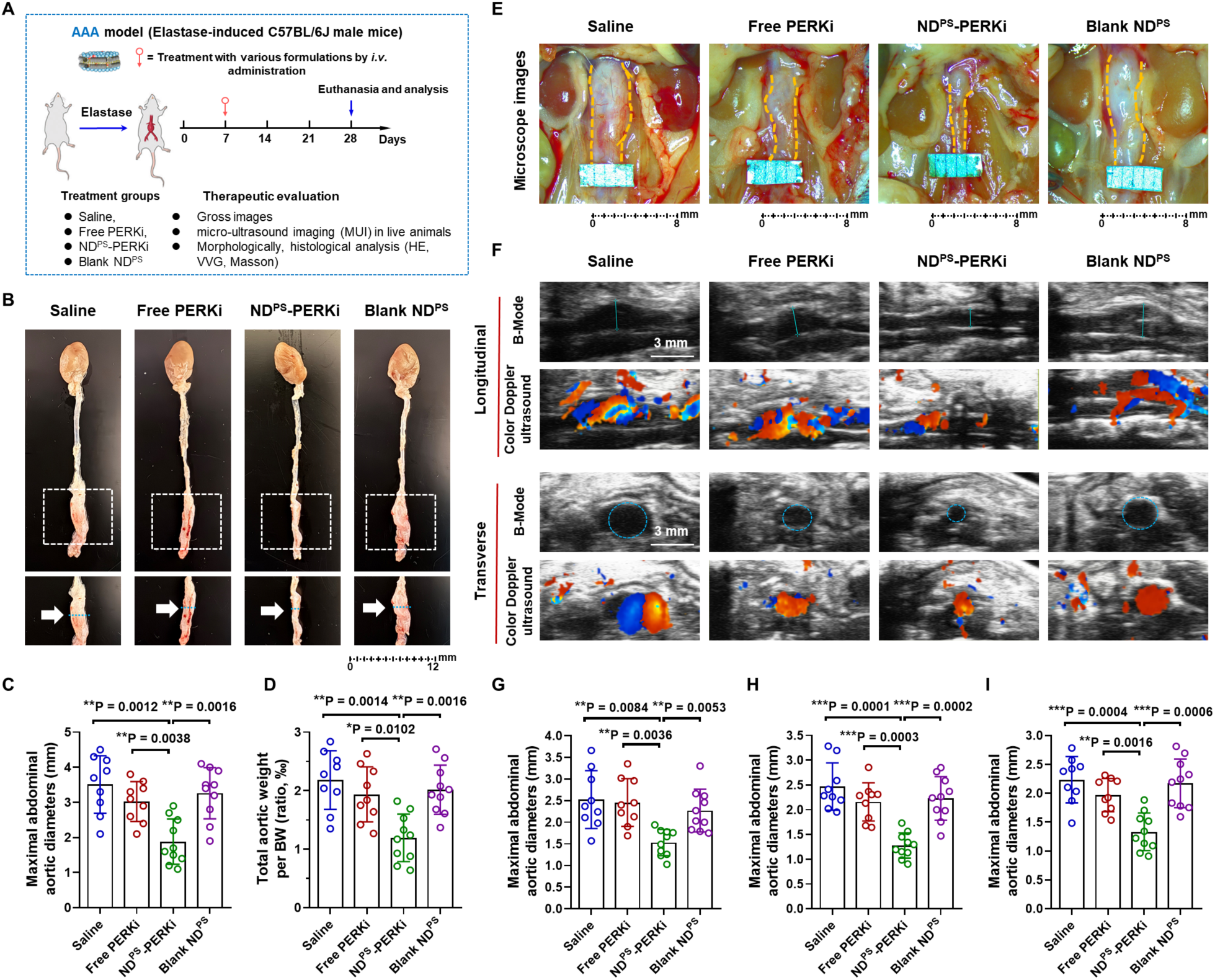
Assessment of therapeutic efficacy of ND^PS^-PERKi in abdominal aortic aneurysm (AAA) mice. **(A)** Schematic illustration of the timeline and treatment protocol for the in vivo study. **(B)** Representative gross images of the whole aortas harvested from AAA mice after different treatments. **(C-D)** Quantification of the maximal diameter of infrarenal abdominal aortas measured using a digital Vernier caliper (C) and the total aortic weight normalized to body weight (BW) in the indicated groups (D) shown in (B) (n = 9 or 10 biologically independent mice). **(E-F)** Representative images of abdominal aortas visualized by stereomicroscope imaging (E) and micro-ultrasound imaging (MUI) in longitudinal and transverse B-mode as well as color Doppler mode (F), from live AAA mice after different treatments. In panel E, yellow dotted lines indicate the injured infrarenal abdominal aortic region. In panel F, blue dotted lines denote the maximal aneurysmal diameter measured in longitudinal and transverse B-mode images. **(G-I)** Quantification of the maximal diameter of infrarenal abdominal aortas measured by stereomicroscope imaging (G) as shown in panel E, and by MUI using the B-mode in longitudinal (H) and transverse (I) as shown in panel F (n = 9 or 10 biologically independent mice). Data were analyzed using one-way ANOVA followed by a Games–Howell post hoc test and are presented as mean ± S.D *P < 0.05, **P < 0.01, ***P < 0.001 and ****P < 0.0001. The cartoon mouse in (A) was created with BioRender.com.

Gross examination of harvested aortas revealed pronounced infrarenal dilation in saline-treated AAA mice, whereas ND^PS^-PERKi-treated mice exhibited attenuated aneurysmal expansion (Fig. 5B). Quantitative analysis confirmed that ND^PS^-PERKi reduced maximal infrarenal aortic diameter by approximately 2-fold as compared to saline, free PERKi, and blank ND^PS^ groups (Fig. 5C). Consistent with this, normalization of aortic weight to body weight demonstrated a significant 45% reduction in aneurysm burden in ND^PS^-PERKi-treated mice compared to saline (Fig. 5D), indicating effective suppression of pathological aortic remodeling.

Aneurysm progression was further assessed longitudinally by stereomicroscopy and micro-ultrasound imaging (MUI). Representative stereomicroscopic images confirmed substantially reduced aortic dilation in the ND^PS^-PERKi group relative to all control treatments (Fig. 5E), corroborated by longitudinal and transverse B-mode MUI and color Doppler imaging (Fig. 5F). Quantitative measurements from stereomicroscopy (Fig. 5G) and MUI in longitudinal (Fig. 5H) and transverse planes (Fig. 5I) showed that ND^PS^-PERKi restrained aneurysmal expansion by almost 2-fold than free PERKi and blank ND^PS^, underscoring its superior in vivo therapeutic efficacy.

These convergent findings demonstrate that ND^PS^-mediated delivery of PERKi effectively suppresses AAA progression, outperforming unencapsulated PERKi and vehicle controls. The superior therapeutic benefit of ND^PS^-PERKi aligns with its enhanced aneurysmal lesion and macrophage targeting observed above, supporting ND^PS^ as a robust nanotherapeutic platform for AAA treatment.

### Preservation of aortic wall structure and matrix integrity by ND^PS^-PERKi

We next performed histological analyses to evaluate whether ND^PS^-PERKi preserves aortic wall integrity and attenuates extracellular matrix remodeling following treatment (Fig. 6). Hematoxylin and eosin (H&E) staining revealed drastic structural disruption in saline-treated AAA mice, whereas ND^PS^-PERKi treatment largely preserved overall aortic wall architecture (Fig. 6A). Masson’s trichrome staining further demonstrated substantial collagen accumulation and disorganized matrix remodeling in control AAA mice, both of which were markedly attenuated by ND^PS^-PERKi treatment (Fig. 6B). Quantitative analysis confirmed a significant reduction in collagen-positive area and collagen fraction in the ND^PS^-PERKi group relative to all other treatment groups (Fig. 6D and E), indicating effective mitigation of pathological fibrotic remodeling.

**Fig. 6.**
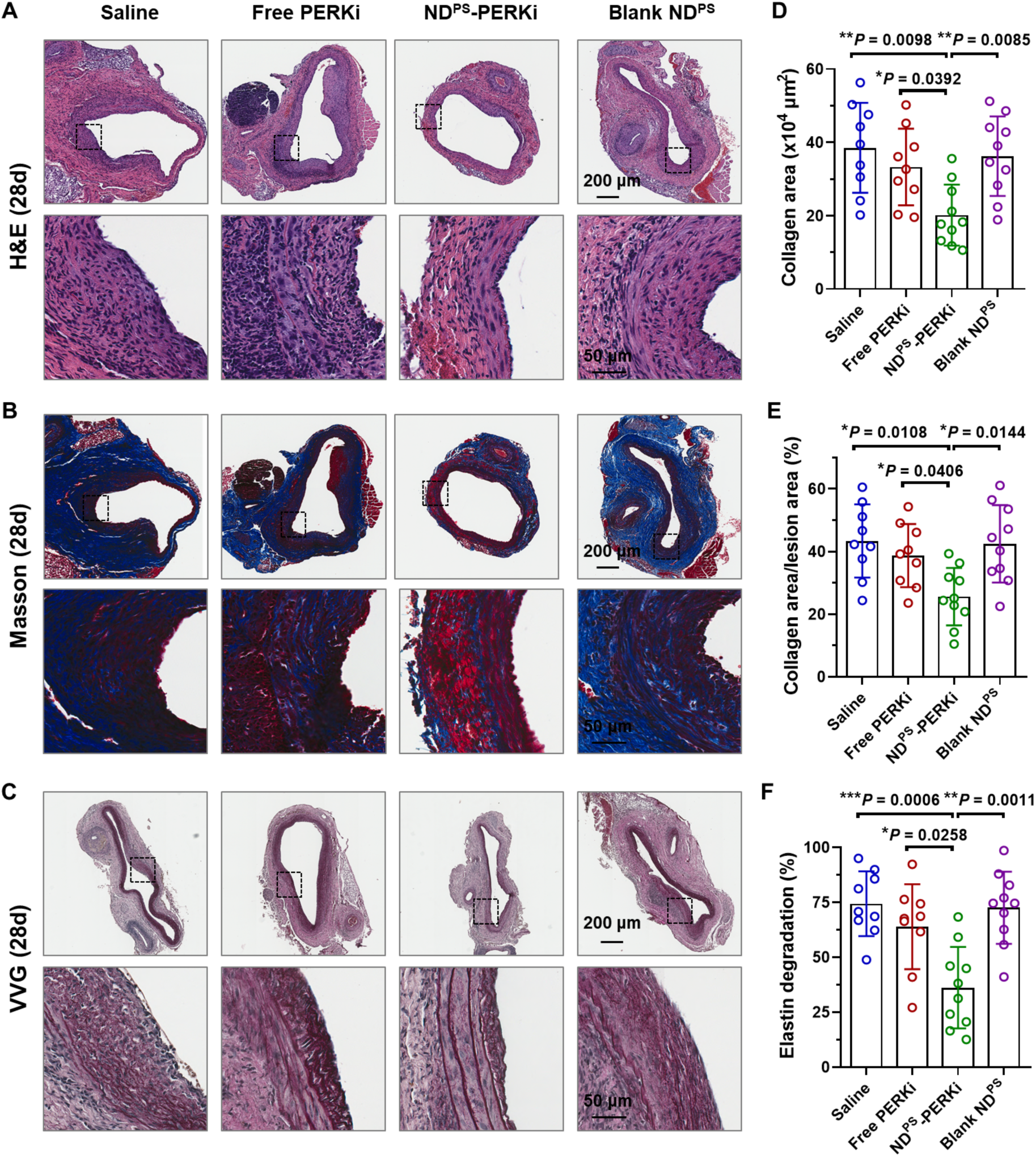
Histological assessment of aortic wall structure and extracellular matrix integrity in AAA mice after ND^PS^-PERKi treatment. **(A-C)** Representative microscopy images of abdominal aortic sections from AAA mice after different treatments stained with (A) hematoxylin and eosin (H&E), (B) Masson’s trichrome, and (C) Verhoeff–Van Gieson (VVG). Black dashed boxes indicate the regions shown in the corresponding magnified images below. Scale bars, 200 μm (top) and 50 μm (bottom). In Masson’s trichrome staining, blue coloration indicates collagen deposition and extracellular matrix remodeling. In VVG staining, elastic lamellae within the aortic wall are visualized, with fragmentation and disruption reflecting elastin degradation characteristic of AAA pathology. **(D-F)** Quantitative analysis of collagen-positive area (D) and relative collagen-positive area (E) measured from Masson’s trichrome staining (B), and elastin fragmentation index (F) calculated from VVG staining (C) in abdominal aortic sections from AAA mice. (n = 9 or 10 biologically independent mice). Data were analyzed using one-way ANOVA with a Games-Howell’s post hoc test, and are shown as mean ± S.D. *P < 0.05, **P < 0.01, ***P < 0.001 and ****P < 0.0001.

Moreover, Verhoeff-Van Gieson (VVG) staining revealed extensive fragmentation and degradation of elastic lamellae in control AAA mice, a hallmark of AAA pathology (Fig. 6C). ND^PS^-PERKi treatment greatly stablized elastin integrity, as evidenced by more continuous elastic lamellae and reduced structural disruption. Quantification of the elastin degradation index confirmed that ND^PS^-PERKi substantially reduced elastin fragmentation compared with unencapsulated PERKi and blank ND^PS^ (Fig. 6F). Collectively, these histological findings demonstrate that ND^PS^-PERKi preserves aortic wall structure and extracellular matrix integrity by attenuating pathological collagen remodeling and elastin degradation, consistent with the marked inhibition of aneurysmal expansion observed in vivo and further supporting therapeutic applications of ND^PS^-PERKi for AAA treatment.

### Mechanistic analysis of the anti-AAA effects of ND^PS^-PERKi in vivo

Having established the therapeutic efficacy of ND^PS^-PERKi, we sought to determine whether treatment modulates macrophage phenotype and attenuates inflammatory responses within aneurysmal lesions (Fig. 7). Given the central role of macrophage-driven inflammation in AAA progression, we first assessed the phenotypic distribution of lesional macrophages across treatment groups.

**Fig. 7.**
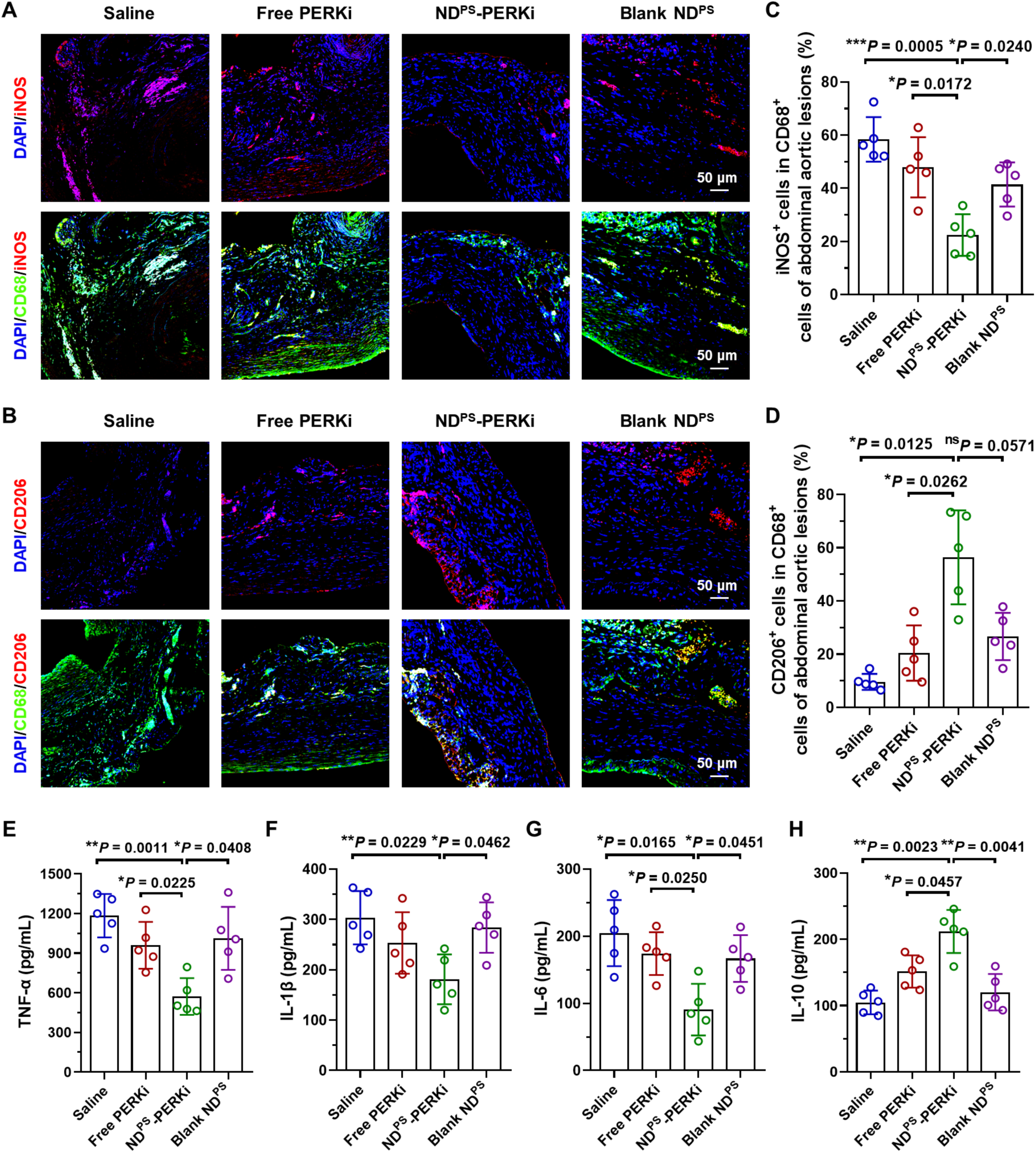
Mechanistic analysis of the anti-AAA effects of ND^PS^-PERKi *in vivo*. (**A-B**) Representative immunofluorescence images showing M1-like (A; iNOS^+^) and M2-like (B; CD206^+^) macrophages among lesional macrophages (CD68^+^) in cross-sections of abdominal aortas from AAA mice after different treatments, including saline, free PERKi, ND^PS^-PERKi, and empty ND^PS^. Blue indicates DAPI staining of nuclei. Yellow signals represent the overlap of green fluorescence (CD68^+^) and red fluorescence (iNOS^+^ or CD206^+^). Figure S4 and S5 present the corresponding original unmerged images. Scale bars, 50 µm. (**C-D**) Quantification of iNOS^+^ (C) and CD206^+^ cells (D; red) within lesional macrophages (green) in cross-sections of abdominal aortas from AAA mice after different treatments (*n* = 5 biologically independent mice, mean ± S.D.). (**E-H**) Serum concentrations of pro-inflammatory cytokines (TNF-α, IL-1β, and IL-6; E-G) and the anti-inflammatory cytokine IL-10 (H) in AAA mice receiving different treatments (*n* = 5 biologically independent mice, mean ± S.D.). Data were analyzed using one-way ANOVA followed by a Games–Howell post hoc test and are presented as mean ± S.D *P < 0.05, **P < 0.01, and ***P < 0.001.

Immunofluorescence staining of aortic sections revealed the highest CD68+iNOS+ signal intensity, indicative of M1-like pro-inflammatory macrophages, in saline-treated AAA mice (Fig. 7A). ND^PS^-PERKi treatment reduced the proportion of M1-like macrophages within lesional areas by 2.6-fold (Fig. 7A, 7C, and Fig. S4), while notably increasing the proportion of anti-inflammatory M2-like macrophages (CD68+CD206+) compared with saline, unencapsulated PERKi, and blank ND^PS^ controls (Fig. 7B, 7D, and Fig. S4). These findings indicate that ND^PS^-PERKi effectively directs lesional macrophages toward an anti-inflammatory M2-like phenotype within the AAA microenvironment.

Complementing these macrophage phenotyping data, we evaluated the systemic anti-inflammatory consequences of macrophage repolarization by quantifying pro-inflammatory cytokines (TNF-α, IL-1β, and IL-6) and the anti-inflammatory cytokine IL-10 in serum from treated AAA mice. ND^PS^-PERKi treatment reduced TNF-α, IL-1β, and IL-6 levels by 30-55% while increasing IL-10 approximately 1.4- to −2-fold relative to all other groups (Fig. 7E-H), consistent with our in vitro observations (Fig. 3G-J). H_2_O_2_ levels were also decreased in the ND^PS^-PERKi group, indicating effective attenuation of systemic oxidative stress (Fig. S6). In line with our cell-based experiments (Fig. 3), these results support that ND^PS^-PERKi exerts simultaneous anti-inflammatory and antioxidative effects in vivo, contributing to robust therapeutic efficacy against AAA progression.

Moreover, these findings indicate macrophage-centered immune reprogramming and redox modulation as the central mechanisms for ND^PS^-PERKi efficacy, wherein coordinated reduction of M1-like inflammation, promotion of M2-like polarization, and suppression of oxidative stress ultimately drive the preservation of aortic wall integrity and restraint of aneurysmal expansion.

### In vivo safety evaluation of ND^PS^-PERKi treatment

Finally, we evaluated systemic toxicity in AAA mice following administration to assess the translational potential of ND^PS^-PERKi (Fig. 8). No significant body weight loss or abnormal fluctuations were observed in ND^PS^-PERKi-treated mice compared with saline, unencapsulated PERKi, or blank ND^PS^ controls throughout the treatment period (Fig. 8A), indicating good overall safety profiles.

**Fig. 8.**
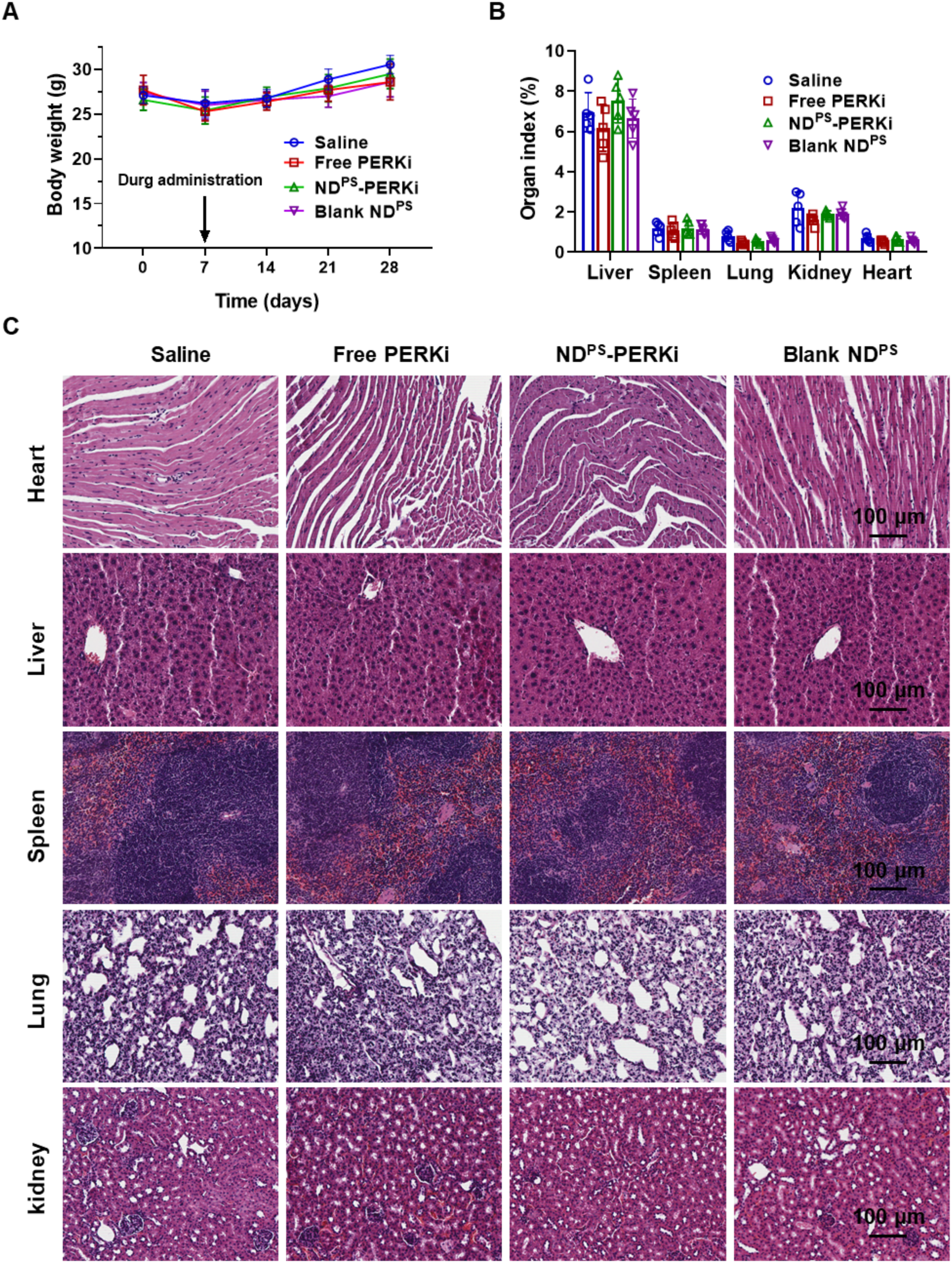
Safety evaluation of AAA mice after ND^PS^-PERKi treatment *in vivo*. **(A)** Body weight of AAA mice across treatment groups over the study period (n = 9 or 10 biologically independent mice). **(B-C)** Organ index of liver, spleen, lung, kidney, and heart (B; n = 5 biologically independent mice), and representative H&E-stained histological sections of major organs harvested from AAA mice following 28 days of various treatments (C). Organ index is defined as: (mass of organ/mass of mouse) ×100%. Scale bar = 100 µm. Data were shown as mean ± S.D.

Consistent with these observations, quantitative analysis of organ indices, including liver, spleen, lung, kidney, and heart, revealed no significant differences among treatment groups (Fig. 8B), suggesting that ND^PS^-PERKi administration does not induce organ swelling, atrophy, or abnormal systemic burden. Histopathological examination of major organs harvested 28 days post-treatment by hematoxylin and eosin (H&E) staining revealed no evident pathological abnormalities, including inflammatory infiltration, tissue necrosis, or structural damage, in any organ of ND^PS^-PERKi-treated mice relative to control groups (Fig. 8C). Serum profiling analysis showed minimal differences in alanine aminotransferase (ALT), aspartate aminotransferase (AST), blood urea nitrogen (BUN), and creatinine (CRE) levels across groups, indicating preserved liver and kidney function (Fig. S7). Hematological analysis further revealed no significant deficiency in red blood cell (RBC), white blood cell (WBC), platelet (PLT), or hemoglobin (HGB) counts following any treatment (Fig. S8).

Therefore, ND^PS^-PERKi exhibited excellent in vivo pharmalogical properties across all parameters examined with no detectable organ damage or systemic toxicity under the therapeutic regimen. Together with its potent aneurysm-suppressive efficacy, these findings support the translational potential of nature-inspired nanodiscs for AAA treatment.

## Discussion

Abdominal Aortic Aneurysm (AAA) remains a life-threatening cardiovascular condition with no approved pharmacological therapy capable of halting its progression.^9–10^ Current clinical management relies predominantly on watchful waiting and surgical intervention, leaving a critical therapeutic gap for patients who are ineligible for or at high risk for operative repair.^6–8^ The present study addresses this unmet clinical challenge by introducing a rationally engineered nanodisc platform (ND^PS^). This platform integrates multi-component targeting with efficient intracellular delivery to enable selective delivery of a potent PERK inhibitor specifically to the macrophage-rich inflammatory microenvironment of aneurysmal lesions. Although PERK has been established as a promising therapeutic target for AAA intervention, effective strategies to selectively engage this pathway remain lacking.^46–49^ While our previous published work utilized a weekly administration to deliver PERKi, our current results demonstrate that targeted nanodisc-assisted PERKi delivery can suppress AAA progression following a single administration in vivo through macrophage-centered immunomodulation. Notably, in contrast to most existing nanotherapeutic and pharmacological approaches that require frequent dosing (e.g., daily oral administration or repeated injections every few days),^17–22^ this platform achieves sustained therapeutic benefit with substantially reduced dosing frequency. This represents a marked improvement over conventional nanoparticle-based strategies, which often suffer from limited lesion specificity and transient efficacy. Together, these findings establish a compelling proof-of-concept for precision nanotherapeutics in vascular disease.

The therapeutic success of ND^PS^-PERKi rests on two synergistic innovations in nanoparticle design. First, the incorporation of an S2P peptide that targets stabilin-2 (STAB2), a receptor upregulated in aneurysmal lesions, enables active homing to diseased aortic sites through recognition of disease-associated receptors^44,45^ on the luminal surface. Second, the inclusion of phosphatidylserine (PS) lipids leverages the well-established recognition of apoptotic cell-surface signals by macrophage scavenger receptors, enabling selective internalization by lesional macrophages. Together, these two targeting mechanisms operate through distinct receptor-mediated pathways, generating a dual-ligand synergy that substantially enhances both lesion accumulation and macrophage uptake relative to single-ligand ND^PC^. Beyond targeting, the use of biomimetic HDL-mimicking nanodiscs represents a critical platform-level innovation. Their structural similarity to endogenous lipoproteins confers low immunogenicity, favorable biocompatibility, and efficient vascular penetration, while previous clinical studies supports their safety profile,^36–39^ providing a translational advantage over many conventional nanomaterials that remain limited by immunogenicity and long-term safety concerns.^40^ Notably, this work represents, to our knowledge, the first application of HDL-mimicking nanodiscs for therapeutic delivery in AAA, highlighting a clinically translatable strategy for vascular-targeted nanomedicine.

Mechanistically, our data provide a coherent picture of how improved intracellular PERKi delivery reshapes the AAA microenvironment. The PERK pathway, a key branch of the unfolded protein response, has emerged as a central regulator of macrophage inflammatory activation and oxidative stress in AAA pathogenesis. By inhibiting PERK signaling, ND^PS^-PERKi effectively scavenged reactive oxygen species and drove macrophage reprogramming from a pro-inflammatory M1-like phenotype toward an inflammation-resolving M2-like state, accompanied by profound reductions in TNF-α, IL-1β, and IL-6 and a significant increase in IL-10. This macrophage-centered immune reprogramming cascaded into tissue-level protection: preserved elastin integrity, reduced pathological collagen remodeling, and attenuated aneurysmal expansion. Mechanistic coherence across molecular, cellular, and structural characterization support that the observed therapeutic benefit reflects genuine biological reprogramming rather than nonspecific effects of nanoparticle administration.

The superior performance of dual-ligand ND^PS^ highlights a broader design principle for nanomedicine: enhancing delivery precision through receptor-synergistic targeting can improve local therapeutic efficacy without increasing systemic exposure. This strategy enables more effective drug accumulation at sites of pathology while minimizing off-target distribution, and may be broadly applicable to other macrophage-driven vascular diseases, including atherosclerosis, myocardial infarction, and stroke, through modular incorporation of disease-specific targeting ligands. From a translational perspective, the ability of ND^PS^-PERKi to achieve therapeutic efficacy following a single administration, together with its favorable safety profile, underscores its potential as a clinically relevant treatment strategy. Moreover, the intrinsic immunomodulatory properties of the nanodisc scaffold suggest that the platform may actively contribute to a pro-resolving therapeutic microenvironment rather than functioning solely as a passive carrier. Collectively, these findings position biomimetic nanodisc-based systems as a promising and versatile platform for precision nanomedicine in vascular and immune-mediated diseases.

### Limitations and Future Directions

Despite these encouraging findings, several limitations warrant further characterizations and point toward productive future directions. The murine elastase-induced AAA model, while widely used and well-validated, does not fully recapitulate the complex pathophysiology of human AAA, which develops over decades and involves distinct genetic, hemodynamic, and immunological factors. Validation in additional preclinical models that more closely approximate human vascular anatomy and disease progression, including porcine or non-human primate systems, will be an important next step toward therapeutic application. Additionally, while this study focused on macrophage phenotype reprogramming and inflammatory suppression as the primary therapeutic mechanism, AAA is a multifactorial disease driven by intricate crosstalk among vascular smooth muscle cells, fibroblasts, T lymphocytes, and neutrophils, in addition to macrophages. Future investigations integrating single-cell or spatial transcriptomic analyses of ND^PS^-PERKi-treated aortic tissue may reveal broader shifts in the vascular immune landscape and uncover additional mechanisms contributing to therapeutic efficacy. Finally, the modular architecture of the ND^PS^ platform presents exciting opportunities for further innovation: alternative therapeutic cargos targeting complementary disease pathways, combination strategies pairing PERKi with anti-inflammatory biologics, or incorporation of stimuli-responsive release mechanisms could further enhance efficacy and versatility. Collectively, these future directions underscore the translational promise of ND^PS^-based nanotherapeutics and the broader potential of precision-engineered nanodisc platforms to transform the treatment landscape for AAA and related macrophage-driven vascular diseases.

### Conclusions

In summary, our study presents a biomimetic nanodisc-based nanotherapeutic platform for AAA treatment, enabling targeted delivery of a PERK inhibitor to lesional macrophages and effective reprogramming of the inflammatory microenvironment. By integrating multi-component targeting with efficient intracellular delivery, this strategy achieves robust therapeutic efficacy following a single administration in vivo. Beyond demonstrating a clinically relevant therapeutic paradigm for AAA, this work highlights the translational potential of nanodisc-based platforms for targeted immunomodulation and establishes a conceptual framework for precision nanomedicine in vascular and other macrophage-associated inflammatory diseases.

## Methods

### Chemicals and Reagents

1,2-dimyristoyl-sn-glycero-3-phosphocholine (DMPC) and 1,2-dioleoyl-sn-glycero-3-phospho-l-serine (DOPS) were purchased from Avanti Polar Lipids. 1,1’-Dioctadecyl-3,3,3’,3’-Tetramethylindodicarbocyanine (DiD) was obtained from ThermoFisher. PERKi (GSK2606414) was purchased from MedChemExpress. Peptides were synthesized by GenScript with >95% purity. All other chemicals were acquired from Sigma.

### Preparation and characterization of nanodiscs

Lipids (100% DMPC or 70%DMPC/30%DOPS at 20 mg/ml) were mixed with DiD (0.1 mg/ml) or PERKi (2mg/ml) and dried under a gentle stream of nitrogen and further with vacuum for 2 h followed by rehydration in reconstitution buffer (25 mM Tris-HCl, pH 7.5, 100 mM NaCl, 1mM DTT).^53,54^ Samples were then extruded through 200 nm filter to generate small unilamellar vesicles (SUVs). Nanodiscs were prepared by incubating SUVs with DeFrMSP-S2P peptides at a ratio of 1/30 at 4°C for 2h with gentle shaking and subjected to centrifugation through 0.22µM filter. Formation of nanodiscs was validated by dynamic light scattering using a DynaPro Nanostar II instrument^25,55^ .

### Negative stain electron microscopy

Formvar/carbon-coated copper grids (01754-F, Ted Pella, Inc.) were glow discharged (15 mA, 25 secs) using PELCO easiGlowTM (Ted Pella, Inc). Nanodiscs (10 µg/ml) were applied onto the grids for 30 secs, followed by staining with 0.75% uranyl formate for 1 minute. Images were collected using a ThermoFisher Science Tecnai G2 TEM (100 kV) equipped with a Veleta CCD camera (Olympus). All TEM data were analyzed using Fiji to determine ND sizes^56^.

### Cell culture

The murine macrophage cell line RAW264.7 was maintained in T-75 culture flasks with vented caps in Dulbecco’s Modified Eagle Medium (DMEM) supplemented with 10% fetal bovine serum (FBS) and 1% penicillin–streptomycin (100 U/mL penicillin and 100 μg/mL streptomycin). Cells were incubated at 37°C in a humidified atmosphere containing 5% CO_2_. The culture medium was refreshed daily, and cells were passaged every other day using 0.25% trypsin–EDTA when they reached the appropriate confluence.

### *In vitro* cellular uptake assay

The cellular uptake of different DiD-labeled sHDL nanoparticle formulations was evaluated by confocal laser scanning microscopy (CLSM) and flow cytometry. RAW264.7 cells were seeded onto glass-bottom dishes at a density of 1 × 10^4^ cells per dish and allowed to adhere overnight. After attachment, the cells were incubated in fresh culture medium supplemented with lipopolysaccharide (LPS, 500 ng/mL) for 12 h. Subsequently, the cells were treated with free DiD or DiD-labeled sHDL-A or sHDL-B at an equivalent DiD concentration of 20 nM for an additional 4 h. After incubation, the cells were washed three times with phosphate-buffered saline (PBS) and stained with DAPI for 15 min. The cells were then washed again with PBS three times and imaged using a confocal laser scanning microscope (Carl Zeiss LSM880) with excitation at 633 nm. For flow cytometry analysis, cells were treated following the same protocol, harvested, re-suspended in PBS, and analyzed using a fluorescence-activated cell sorting system (Accuri C6, BD Biosciences) with an excitation wavelength of 633 nm. Flow cytometry data were analyzed using FlowJo software, and quantitative analyses were performed using GraphPad Prism 8.

### In vitro ROS scavenging capability assay

The intracellular reactive oxygen species (ROS) scavenging capability of different formulations was evaluated using the 2′,7′-dichlorofluorescin diacetate (DCFH-DA; Thermo Fisher Scientific, catalog no. D399) probe according to the manufacturer’s instructions. RAW264.7 cells were seeded into 12-well plates at a density of 1.0 × 10^5^ cells per well and incubated for 12 h to allow cell attachment. The culture medium was then removed, and the cells were pretreated with either fresh medium alone (model control) or different formulations, including free PERK inhibitor (PERKi), ND^PC^-PERKi, or ND^PS^-PERKi, at an equivalent PERKi concentration of 20 μg/mL for 2 h. Subsequently, the cells were co-incubated with LPS (500 ng/mL) for 12 h to induce oxidative stress or inflammation. After stimulation, the cells were incubated with fresh medium containing DCFH-DA (20 μM) at 37°C for 30 min. The cells were then washed three times with phosphate-buffered saline (PBS), and the fluorescence intensity of intracellular DCF was quantified by flow cytometry (BD Biosciences, San Jose, CA, USA) at an excitation/emission wavelength of 488/525 nm. Flow cytometry data were analyzed using FlowJo software, and quantitative analyses were performed using GraphPad Prism 8.

### Measurement of pro-inflammatory and anti-inflammatory cytokine levels

The secretion levels of pro-inflammatory and anti-inflammatory cytokines were quantified by enzyme-linked immunosorbent assay (ELISA). RAW264.7 cells were treated following the same experimental protocol as described in the “*In vitro* ROS scavenging capability assay*”*. Briefly, cells were pretreated with either culture medium alone (model control) or different formulations, including free PERK inhibitor (PERKi), ND^PC^-PERKi, or ND^PS^-PERKi, at an equivalent PERKi concentration of 10 μg/mL for 2 h, followed by co-incubation with lipopolysaccharide (LPS, 500 ng/mL) for 12 h. After treatment, the culture supernatants were collected and centrifuged to remove cellular debris. The secretion levels of representative pro-inflammatory cytokines, including tumor necrosis factor- α (TNF-α), interleukin-1β (IL-1β), and interleukin-6 (IL-6), as well as the anti-inflammatory cytokine interleukin-10 (IL-10), were determined using commercially available ELISA kits according to the manufacturers’ instructions. Absorbance was measured using a microplate reader, and cytokine concentrations were calculated based on standard curves.

### Animals and Abdominal Aortic Aneurysm (AAA) model

All animal studies were conducted under the National Institutes of Health Guide for the Care and Use of Laboratory Animals. Research protocols are approved by the Institutional Animal Care and Use Committees and Institutional Biosafety Committee at University of Virginia. 10-12 weeks male C57BL/6J mice were purchased from the Jackson Laboratory and maintained in a temperature and humidity-controlled animal facility under a 12 h light-dark cycle.

A modified elastase procedure was performed based on previously published method^46^. Mice were subject to drinking water modification with 0.2% β-Aminopropionitrile (BAPN) from one day prior to the elastase challenge till the day harvest. After anesthesia by inhaled isoflurane and sterile preparation of the incision sites, a midline incision was made and the infrarenal region of the abdominal aorta was exposed. The mice then received topical application of 0.6 U porcine pancreatic elastase for 8 min onto the abdominal aortic segment. After elastase incubation, the elastase was removed and the residual elastase solutions were washed by saline at least 3 times. Bowel contents were returned to the original positions, and the muscle layer and skin layer were closed by 4-0 suture. On day 7 after elastase-induced AAA surgery, mice received tail vein injections of GSK2656157 (a selective protein kinase R–like endoplasmic reticulum kinase PERK inhibitor; GSK2606414)–loaded nanodiscs at a dose of 5 mg/kg, or the corresponding control treatments. High-resolution ultrasound imaging was performed using a Vevo 2100 system (FUJIFILM VisualSonics) to acquire abdominal aortic images one day prior to the experimental endpoint. On day 28, mice were anesthetized and the abdominal aortas were carefully dissected for measurement of maximal aortic diameter. Subsequently, abdominal aortas and major organs, including the heart, liver, lung, spleen, and kidney, were collected following systemic perfusion with phosphate-buffered saline (PBS) and 4% paraformaldehyde (PFA) for further histological analysis.

### Echocardiography

To assess changes in aortic diameter, high-resolution ultrasound imaging was performed using a Vevo 2100 system (FUJIFILM VisualSonics). Mice were anesthetized with 1.2–1.5 vol% isoflurane, and the heart rate was maintained at approximately 400 beats per minute during image acquisition. The maximal diameter of the infrarenal abdominal aorta was measured from the ultrasound images. An aneurysm was defined as a ≥50% increase in infrarenal aortic diameter relative to tthe diameter at day 0 pre-challenge.

### *In vivo* biodistribution and AAA-targeting capability assessment

To evaluate the in vivo biodistribution of different formulations, elastase-induced abdominal aortic aneurysm (AAA) model mice or healthy C57BL/6J mice were intravenously administered with 100 μL of ND^PC^-DiD or ND^PS^-DiD at an equivalent DiD dose of 2 μM per mouse, at day 7 post the surgery. At 24 h post-injection, the mice were euthanized and perfused with phosphate-buffered saline (PBS) to remove residual circulating DiD. The aortas and major organs, including the hearts, livers, spleens, lungs, and kidneys, were harvested and subjected to *ex vivo* near-infrared fluorescence (NIRF) imaging using an IVIS Lumina imaging system (PerkinElmer). Fluorescence signals were acquired at excitation and emission wavelengths of 633 and 670 nm, respectively, and quantitative analysis was performed using the manufacturer’s software.

To further assess macrophage-targeting capability in the abdominal aorta of AAA mice, abdominal aortic tissues were collected, fixed in ice-cold acetone, embedded, and sectioned at a thickness of 5 μm. The tissue sections were incubated overnight at 4 °C with an anti-CD68 primary antibody (Abcam, ab283654, rabbit, 2 μg/mL). After washing, the sections were incubated with an Alexa Fluor® 488–conjugated goat anti-rabbit IgG secondary antibody (Abcam, ab150077, 2 μg/mL) for 2 h at room temperature in the dark. Nuclei were subsequently counterstained with DAPI. Finally, the stained sections were imaged using a confocal laser scanning microscope (Carl Zeiss LSM880), and fluorescence signal quantification was performed using ImageJ software.

### *In vivo* therapeutic evaluation of sHDL formulations in AAA mice

The therapeutic efficacy of sHDL formulations was evaluated in an elastase-induced abdominal aortic aneurysm (AAA) mouse model. AAA was induced in C57BL/6J mice by topical elastase challenge as described above. After model establishment, mice were randomly assigned to different treatment groups (*n* = 10 per group), including Saline, free PERK inhibitor (PERKi), ND^PS^-PERKi, and blank ND^PS^ (without loading PERKi). Therapeutic intervention was initiated on day 7 post-AAA induction. Mice received a single intravenous injection via the tail vein of the indicated formulations on day 7. The dose of PERKi was 5 mg/kg per mouse, and the injection volume was kept constant across all groups. This dose was selected based on our prior studies and optimization experiments. For sHDL-based formulations, the administered dose was normalized to an equivalent amount of PERKi. Mice were monitored throughout the study for changes in body weight and general health status. At the experimental endpoint (day 28 post-induction), mice were euthanized and perfused with saline. The abdominal aortas were carefully harvested for therapeutic evaluation. The maximal abdominal aortic diameter was measured using multiple complementary approaches, including digital calipers, stereomicroscopy, and high-frequency ultrasound imaging, to ensure accurate and reliable assessment. Following diameter measurement, aortic tissues were further processed for histopathological analysis.

### Histological Analysis

Abdominal aortas were harvested, perfused with saline, and fixed with 4% paraformaldehyde, followed by embedding in paraffin and serial sectioning. Paraffin-embedded sections (5 μm thick) were prepared for histological analysis. Hematoxylin and eosin (H&E) staining was performed to evaluate general tissue morphology, and Masson’s trichrome staining was conducted to assess collagen deposition. These stainings were carried out by the Histology Core Facility of the Robert M. Berne Cardiovascular Research Center at the University of Virginia. Verhoeff–Van Gieson (VVG) staining was performed using a commercial staining kit (Sigma, 115974) according to the manufacturer’s instructions to evaluate elastin degradation. Elastin degradation was quantified as the percentage of degraded elastic fiber length relative to the total internal elastic lamina length. Collagen content was quantified from Masson’s trichrome–stained sections using ImageJ software.

### Immunofluorescence Staining

Fluorescent immunostaining was performed following our previously published protocol. Paraffin-embedded arterial sections were deparaffinized and subjected to antigen retrieval using antigen-retrieval buffer (ab93678, Abcam). After washing with PBS, the sections were permeabilized with PBS containing 0.25% Triton X-100 for 15 min and washed three times with PBS (10 min each). The sections were then blocked with PBS containing 5% bovine serum albumin (BSA) for 1 h at room temperature. Subsequently, the tissue sections were incubated overnight with primary antibodies against CD68 (BMA Biomedicals, T-3003, 2 μg/mL), iNOS (Abcam, ab15323, 1 μg/mL), or CD206 (Abcam, ab64693, 1 μg/mL). After washing three times with PBS, the sections were incubated with the corresponding secondary antibodies, including Alexa Fluor 488-conjugated or Alexa Fluor 594-conjugated anti-rabbit/mouse IgG (Invitrogen, A-11001 and A-11037), for 1 h at room temperature. Cell nuclei were counterstained with DAPI mounting medium (P36962, Thermo Fisher Scientific). Finally, the immunostained sections were imaged using a confocal laser scanning microscope (CLSM, Zeiss LSM880, Germany), and fluorescence signals were quantified using ZEN 2.3 software (Blue edition).

### Quantification and statistical analysis

All experiments were performed with at least three independent biological replicates, yielding consistent results. Data normality was assessed using the D’Agostino–Pearson or Shapiro–Wilk test. For comparisons between two groups with normally distributed data, a two-tailed Student’s *t* test was applied. For comparisons among more than two groups, one-way analysis of variance (ANOVA) followed by Tukey, Dunnett’s T3, or Games–Howell post hoc tests was used. When data deviated from normality, pairwise comparisons were performed using the Mann–Whitney U test, whereas multi-group comparisons were conducted using the Kruskal–Wallis test. Statistical significance was defined as *P* < 0.05, with increasing levels of significance indicated as \**P* < 0.05, \*\**P* < 0.01 and \*\*\**P* < 0.001. Data are presented as mean ± SD. All statistical analyses were performed using GraphPad Prism 8.0 (GraphPad Software, USA).

## ASSOCIATED CONTENT

### Supporting Information

The Supporting Information is available free of charge at the ACS Publications website. The incorporation of the targeting peptide improves the uptake efficiency of sHDL; Blank NDs platform itself promotes inflammation resolution; Biodistribution of ND^PC^-DiD or ND^PS^-DiD in other major organs of AAA mice; ND^PS^-PERKi therapy reduces pro-inflammatory macrophage polarization; ND^PS^-PERKi therapy promotes anti-inflammatory macrophage polarization; ND^PS^-PERKi therapy reduces lesional oxidative stress production; Biochemical assays of hepatic and kidney functions; Assays of typical hematological parameters.

### Materials availability

This study did not generate new unique reagents.

### Data and code availability

The data supporting the findings in this study are available within the paper and its supplemental information. All data generated in this study are available from the lead contact on reasonable request.

## Acknowledgment

We thank the Bao lab members for discussion and suggestions. H.B. acknowledge support from NIH (DP2GM140920 and R35GM156801), Virginia Common Health Research Board (CHRB-207-01-25), University of Virginia Comprehensive Cancer Center (P30CA044579) and an institutional research grant from the American Cancer Society. In addition, this research was conducted while H.B. was a Hevolution/AFAR New Investigator Awardee in Aging Biology and Geroscience Research. L.-W.G. acknowledges support from NIH (R01HL167902, R01HL172888).

## Author Contribution

Z.H., Y.H.,, L.G. and H.B. conceived the project and wrote the manuscript with input from all authors. Z.H., and Y.H., carried out in vivo studies of nanodiscs-assisted delivery of PERKi. H.Z. and J.X. performed cell-based experiments. Y.W. and Q.R. contributed reconstitutions and characterizations of nanodiscs by gel electrophoresis, DLS and electron microscopy. Q.W. assisted in confocal microscopy experiments.

## Data availability

All other data supporting the findings of this study are available within this manuscript.

## Supplementary Information

**Fig. S1.**
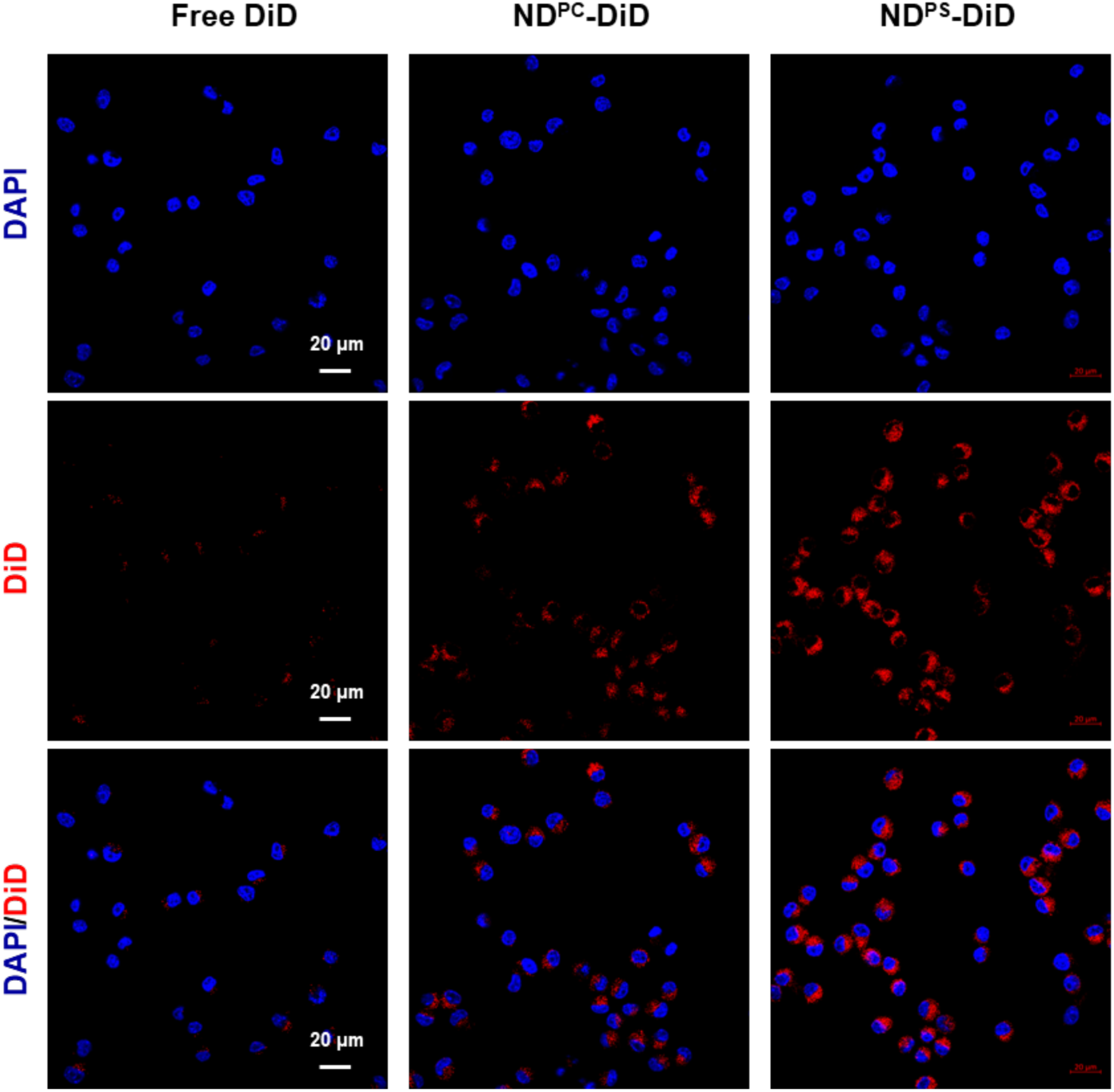
The incorporation of the targeting peptide improves the uptake efficiency of sHDL by RAW264.7 macrophages. Representative confocal microscopy images of RAW264.7 macrophages treated with free DiD and DiD-labeled ND^PC^ or ND^PS^. The cell nuclei were stained with DAPI (blue). The merge images are also presented in Fig. 3A. Scale bars = 20 µm.

**Fig. S2.**
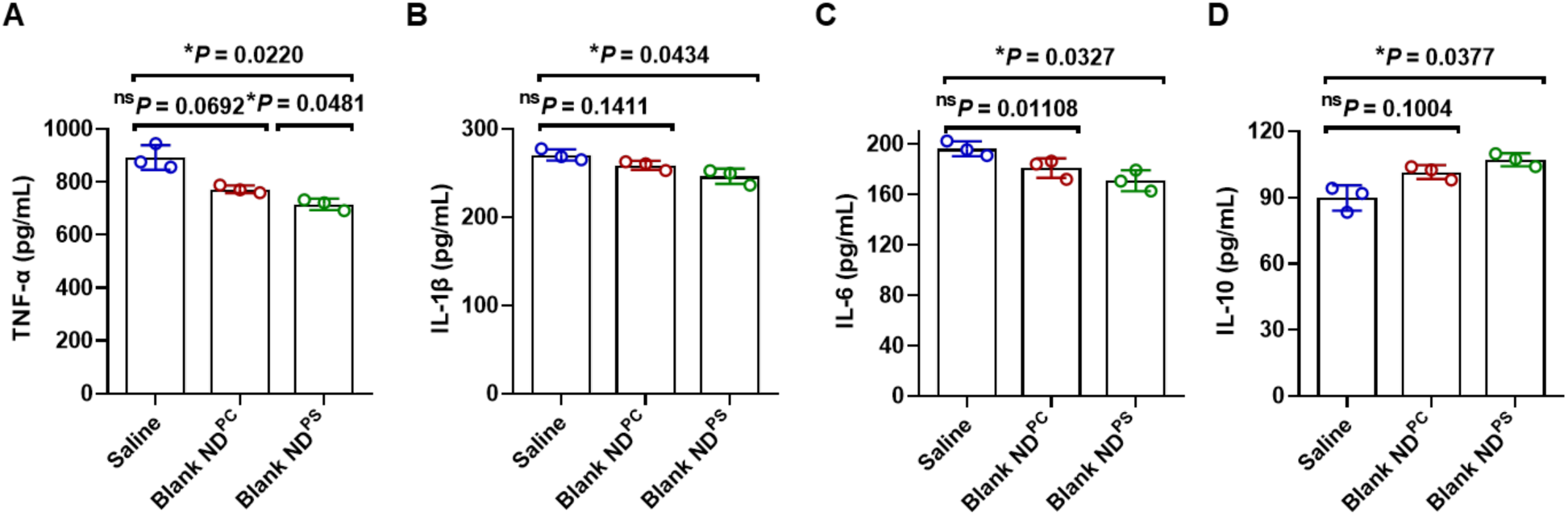
Blank NDs moderately promotes inflammation resolution. (**A-D**) Secretion levels of pro-inflammatory cytokines (TNF, IL-1β and IL-6) (A-C) and anti-inflammatory cytokine IL-10 (D) by RAW264.7 cells were measured by ELISA. Data were analyzed using one-way ANOVA with a Games-Howell post hoc test and shown as mean ± S.D. (n = 3 biologically independent samples). Statistical significance is indicated as*P < 0.05, **P < 0.01, and P > 0.05 denotes no significance.

**Fig. S3.**
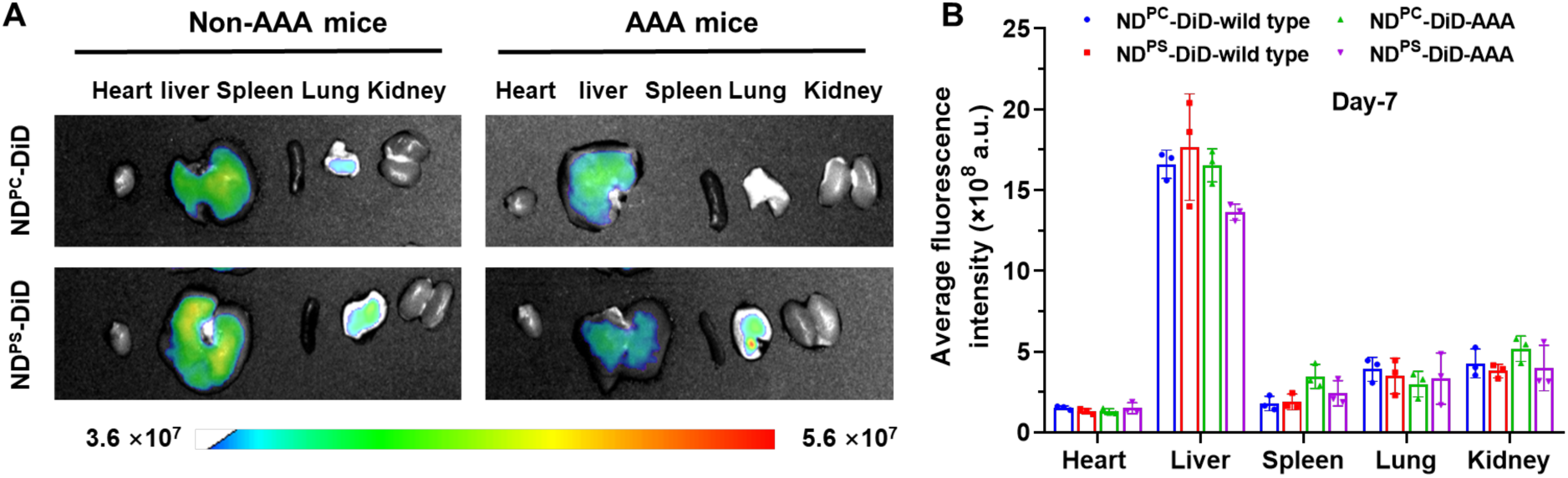
Biodistribution of ND^PC^-DiD or ND^PS^-DiD in other major organs of AAA mice. **(A-B)** Representation of IVIS images (A) and quantification of DiD fluorescence signals (B) in the main organs (heart, liver, spleen, lung, and kidney) 24 h after administration of various formulations in abdominal aortic aneurysm (AAA) mice or C57BL/6J healthy non-AAA mice. AAA mice were generated by incubating ∼0.6U elastase around the abdominal aorta for 8 minutes, followed by a 7-day waiting period before administration of DiD-labelled sHDLs. Data were shown as mean ± S.D. (n = 3 biologically independent mice).

**Fig. S4.**
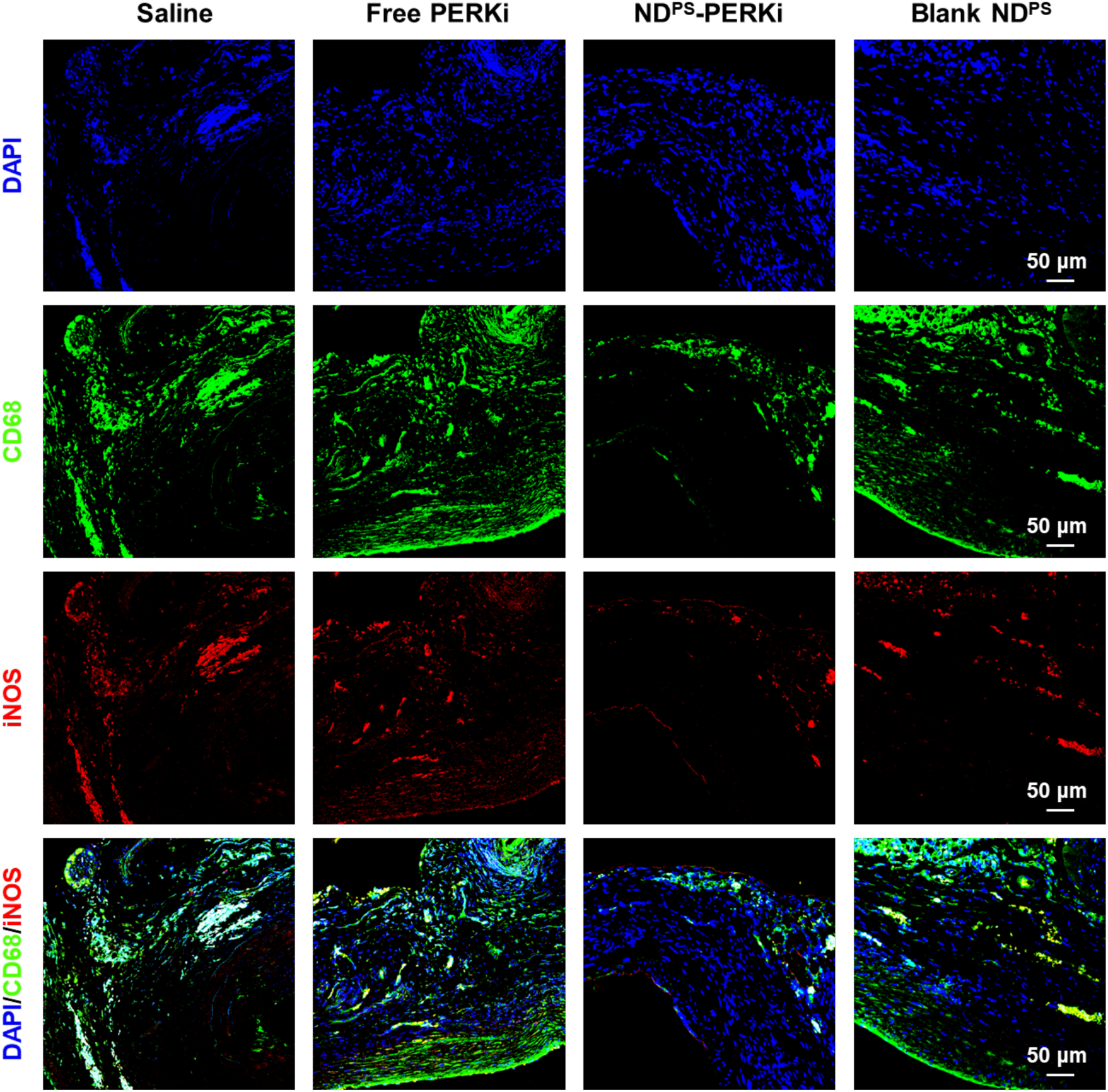
ND^PS^-PERKi therapy reduces pro-inflammatory macrophage polarization. Elastase-induced abdominal aortic aneurysm (AAA) mice were treated with saline, free PERK inhibitor (PERKi), ND^PS^-PERKi, or blank ND^PS^ (without PERKi loading). PERKi was administered at 5 mg/kg per mouse. Immunofluorescence images of abdominal aortic sections depicting iNOS^+^ cells (iNOS, red), macrophages (CD68, green), and nuclei (DAPI, blue) within aneurysmal lesions. The corresponding merged images are also shown in Figure 7A. Scale bars, 50 µm.

**Fig. S5.**
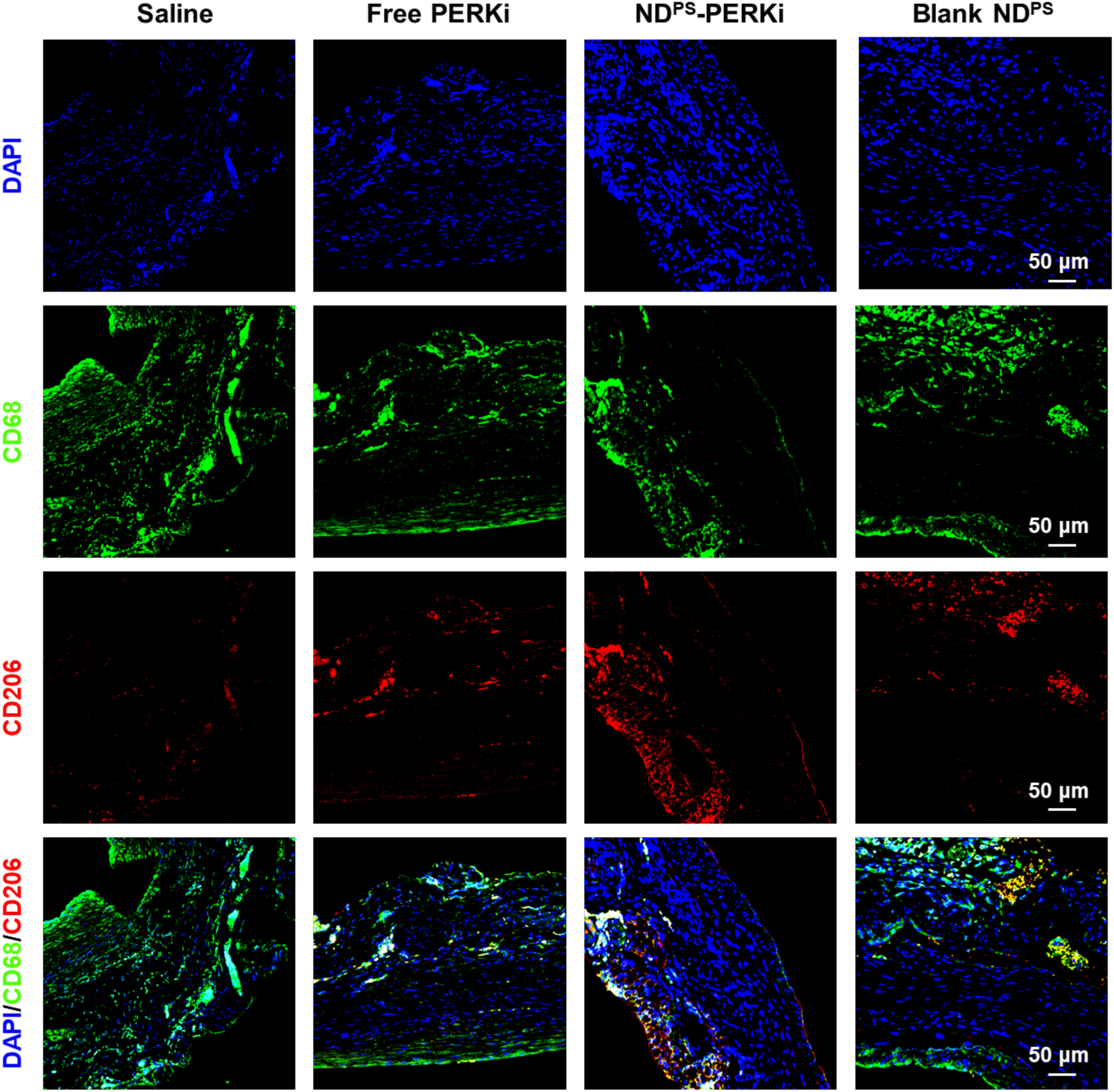
ND^PS^-PERKi therapy promotes anti-inflammatory macrophage polarization. Elastase-induced abdominal aortic aneurysm (AAA) mice were treated with saline, free PERK inhibitor (PERKi), ND^PS^-PERKi, or blank ND^PS^ (without PERKi loading). PERKi was administered at 5 mg/kg per mouse. Immunofluorescence images of abdominal aortic sections depicting CD206^+^ cells (CD206, red), macrophages (CD68, green), and nuclei (DAPI, blue) within aneurysmal lesions. The corresponding merged images are also shown in Figure 7B. Scale bars, 50 µm.

**Fig. S6.**
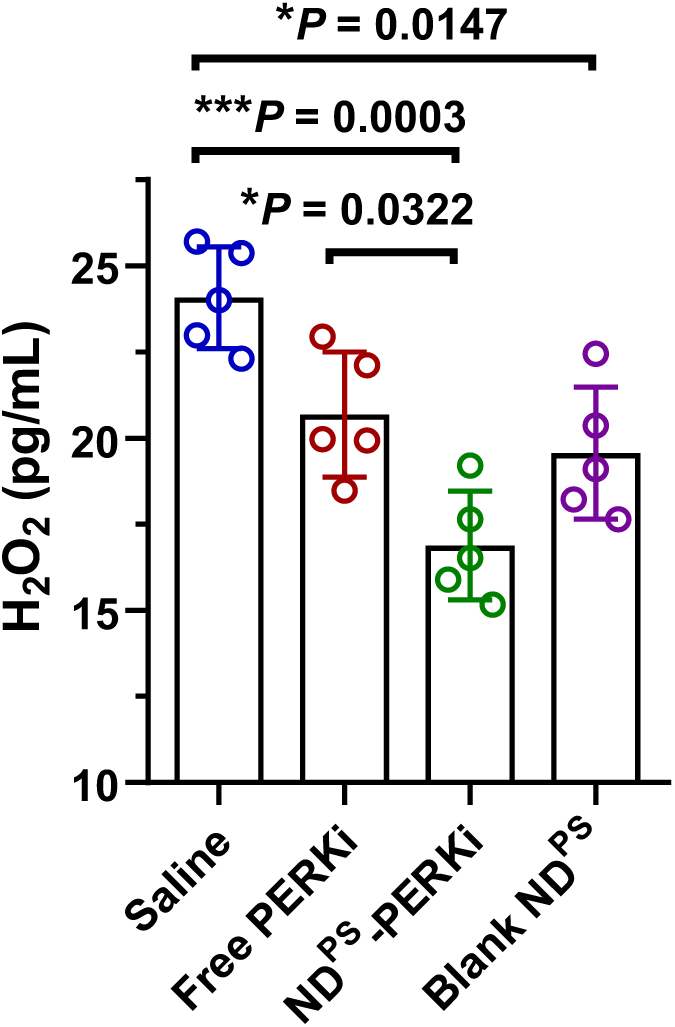
ND^PS^-PERKi treatment reduces lesional oxidative stress production. Elastase-induced abdominal aortic aneurysm (AAA) mice were treated with saline, free PERK inhibitor (PERKi), ND^PS^-PERKi, or blank ND^PS^ (without PERKi loading). PERKi was administered at 5 mg/kg per mouse. Serum concentrations of H_2_O_2_ in AAA mice after receiving different treatments (*n* = 5 biologically independent mice, mean ± S.D.). Data were analyzed using one-way ANOVA followed by a Games–Howell post hoc test and are presented as mean ± S.D *P < 0.05, **P < 0.01, and ***P < 0.001.

**Fig. S7.**
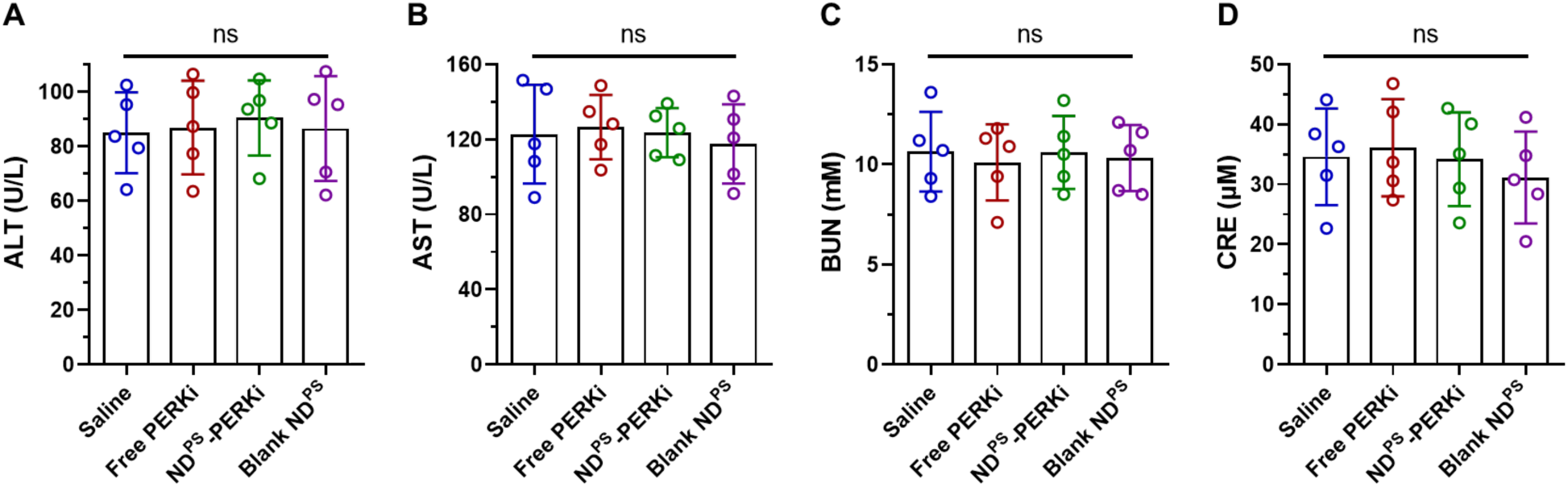
ND^PS^-PERKi treatment has no adverse effect on hepatic and renal function. Hepatic and renal function were characterized by measuring serum levels of (**A**), alanine aminotransferase (ALT), (**B**) aspartate aminotransferase (AST), (**C**) blood urea nitrogen (BUN), and (**D**) creatinine (CRE). Data were presented as mean ± S.D. (n = 5 biologically independent mice) and were analyzed by one-way ANOVA with Tukey post hoc test. No statistically significant difference was observed among different groups.

**Fig. S8.**
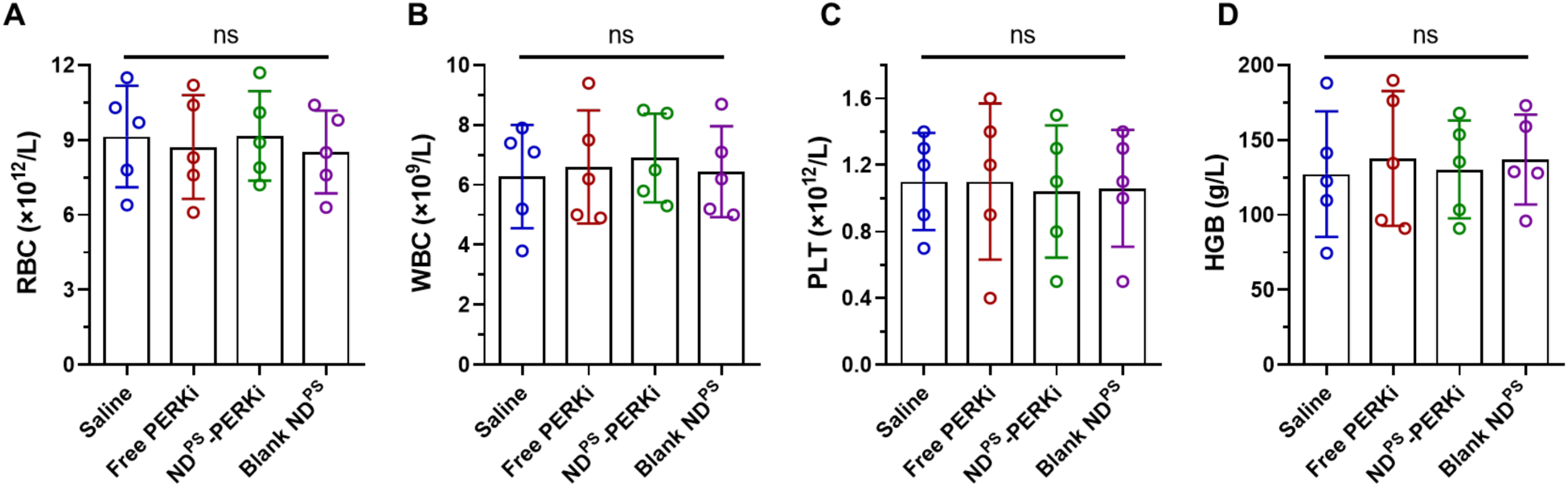
Hematological parameters are unaffected by ND^PS^-PERKi treatment. Hematological parameters were assessed following NDPS-PERKi treatment and control conditions by measuring (**A**) red blood cell (RBC), (**B**) white blood cell (WBC), (**C**) platelet (PLT), and (**D**) hemoglobin (HGB) counts. Data are presented as mean ± S.D. (n = 5 biologically independent mice) and were analyzed by one-way ANOVA with Tukey post hoc test. No statistically significant differences were observed among different groups.

